# Single-Atom Modification Enables Reversal of Degradation Selectivity between NSD2 and GSPT1

**DOI:** 10.64898/2026.09.09.750440

**Authors:** Weizhong Shen, Yihan Liu, Lianchao Liu, Yingfan Liu, Yan Zhang, Yihan Chen, Lei Wang, Zhiyi Zhou, Angela Zhang, Hui Shen, Fengtao Zhou, Weixue Huang, Xiaomei Ren, Abhijit Parolia, Yong Xu, Yang Zhou, Zhen Wang, Arul M. Chinnaiyan, Ke Ding

**Author notes:** **Corresponding Authors:** (Y.Z.); (Z.W.); (A.M.C.); (K.D.). **Author Contributions:** W. S., Y. L., L. L., Y. L., Y. Z., and Y. C. contributed equally to this work.

## Abstract

Both NSD2 and GSPT1 represent compelling targets for anti-cancer drug discovery. Herein, we carried out single-atom modification of **LLC0424**, an NSD2 degrader with GSPT1 neosubstrate engagement, to yield two structurally analogous molecules with divergent modes of action: **424-ND**, an NSD2-selective degrader, and **424-GD**, a GSPT1-selective degrader. **424-ND** drove selective NSD2 degradation and suppressed androgen receptor (AR) signaling in prostate cancer cells. **424-GD** induced selective degradation of GSPT1 and upregulated the integrated stress response markers ATF4 and ATF3. Biolayer interferometry revealed distinct ternary complex cooperativity profiles in the presence of NSD2 or GSPT1, which directly correlate with the observed biased degradation activity. Molecular dynamics and metadynamics simulations showed that the compounds adopt distinct low-energy conformational ensembles with different spatial orientations, likely underlying their divergent cooperativity and selectivity. These findings demonstrate that minimal structural alteration permits precision control over degrader target selectivity, while furnishing selective chemical probes for NSD2 and GSPT1.

## INTRODUCTION

Nuclear receptor–binding SET domain–containing protein 2 (NSD2) is a histone methyltransferase that catalyzes the dimethylation of histone H3 lysine 36 (H3K36me2)^1^. As a functionally important epigenetic regulator, aberrant NSD2 expression, somatic mutations, or chromosomal translocations have been shown to elevate global H3K36me2 levels, leading to epigenomic reprogramming that promotes tumor progression across multiple cancer types^2–6^. However, the biological functions of NSD2 and particularly its relationship with cancer remain incompletely understood, highlighting the need for selective chemical tools. In recent years, while several NSD2 degraders have been reported (Figure 1), including **MS159**^7^, **UNC8732**^8^, **3**^9^, and **ND-L11B**^10^, the proteomic selectivity for most compounds remains unknown. For example, **MS159**, the first-in-class NSD2 degrader, not only exhibited a DC_50_ value of 5.2 μM for NSD2 in 293FT cells but also effectively degraded cereblon (CRBN) neo-substrates IKZF1 and IKZF3. The only series of NSD2 degraders that displayed favorable degradation selectivity for NSD2, exemplified by **UNC8732**, was found to recruit FBXO22 ligase by metabolizing the primary amine to an aldehyde species. Therefore, the development of highly selective NSD2 degraders with distinct mechanisms of action remains essential to enable precise functional studies and to accurately evaluate the therapeutic potential of NSD2 targeting.

**Figure 1.**
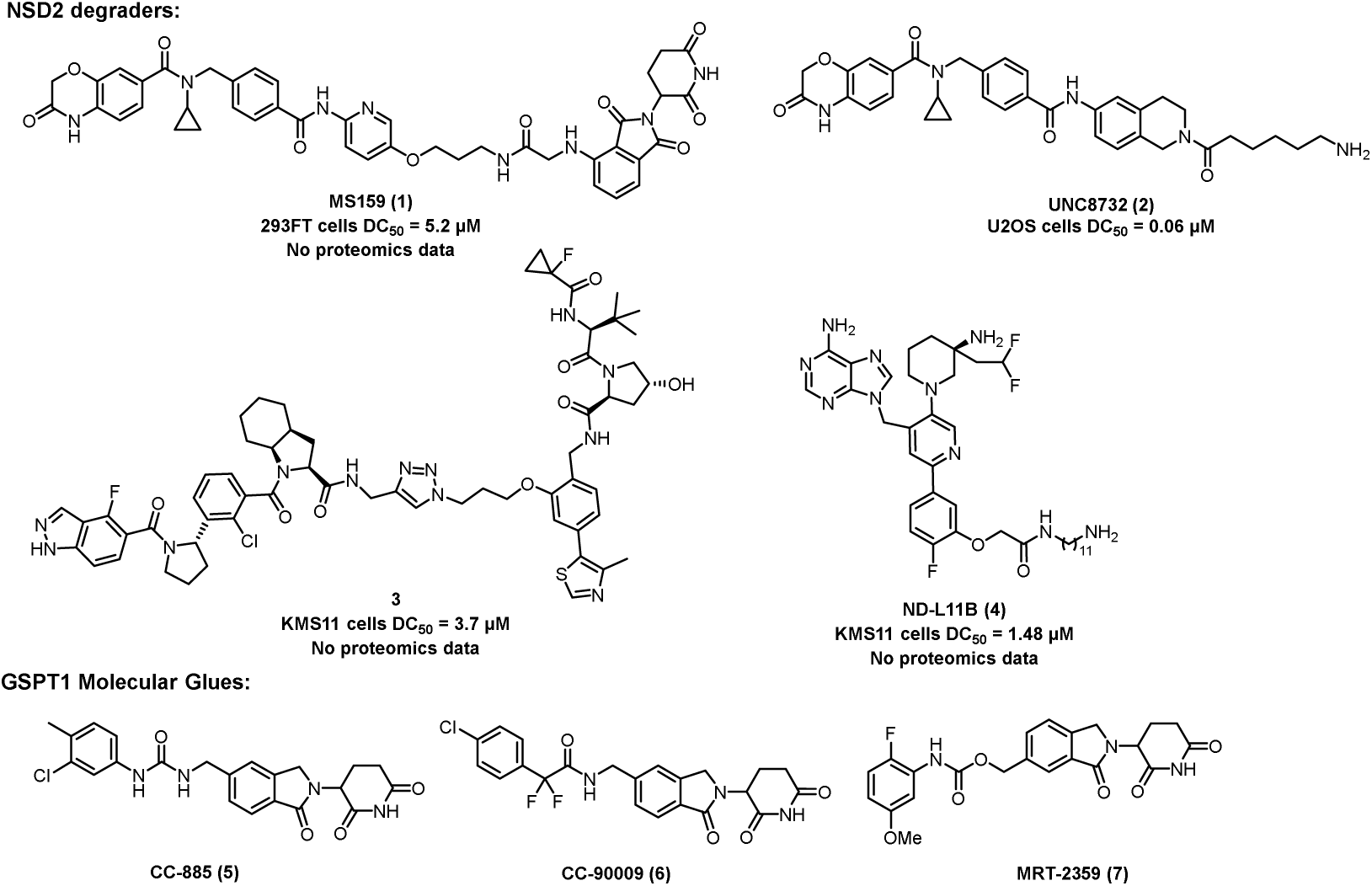
Chemical structures of representative NSD2 degraders and GSPT1 degraders.

G1 to S phase transition protein 1 (GSPT1), also known as eukaryotic peptide chain release factor 3a (eRF3a), is a GTPase that functions as a core component of the translation termination machinery^11, 12^. By cooperating with eukaryotic release factor 1 (eRF1) in a guanosine triphosphate (GTP) –dependent manner, GSPT1 mediates ribosome release at stop codons and is essential for maintaining global protein synthesis and cell viability^13, 14^. Although GSPT1 is required for normal cellular function, studies have shown that cancer cells, particularly hematologic malignancies, are more dependent on GSPT1 activity, suggesting that GSPT1 may serve as a potential therapeutic target^15–17^. As show in Figure 1, **CC-885** was the first reported GSPT1 molecular glue^18^, and subsequent optimization efforts led to the development of **CC-90009**^19^ and **MRT-2359**^20^, both of which advanced into clinical evaluation. However, no GSPT1 degrader has yet been approved for clinical use, and new GSPT1 degraders remain desirable.

Here, we performed single-atom modification of the NSD2 degrader **LLC0424**, which displays unintended neosubstrate engagement of GSPT1, to achieve inversion of degradation selectivity between NSD2 and GSPT1. Specifically, our work revealed that subtle structural modification biased the scaffold toward either an NSD2-selective degrader (**424-ND**) or a GSPT1-selective degrader (**424-GD**). Biophysical analyses indicated that these precise chemical modifications modulate the stability of CRBN-mediated ternary complex formation, thereby inverting degradation selectivity. Molecular dynamics reveals distinct conformational landscapes of the compounds, providing a structural basis for the observed inversion of degradation selectivity.

## RESULTS AND DISCUSSION

### Single-atom modification of LLC0424 yields NSD2-selective degrader (424-ND) and GSPT1-selective degrader (424-GD)

We selected our previously reported degrader **LLC0424** as the starting point for optimization. Despite its faster degradation kinetics toward NSD2 over GSPT1, **LLC0424** exhibited potent dual potency against both targets in VCaP prostate cancer cells, with DC_50_ values of 81 nM and 88 nM, respectively^21^. We chose to do modification of the piperidine linker and the thalidomide because (1) the rigid piperidine linker bridges the NSD2 binder and the E3 ligase ligand and governs ternary complex geometry; subtle change could alter molecular conformation, thereby influencing ternary complex stability/cooperativity and ultimately modulating degradation selectivity; (2) the electrostatic complementarity at the thalidomide–CRBN interface similarly governs ternary complex geometry; fine-tuning of the electronic properties of thalidomide can reshape ternary complex stability/cooperativity, enabling reprogramming of degradation selectivity (Figure 2)^22^.

**Figure 2.**
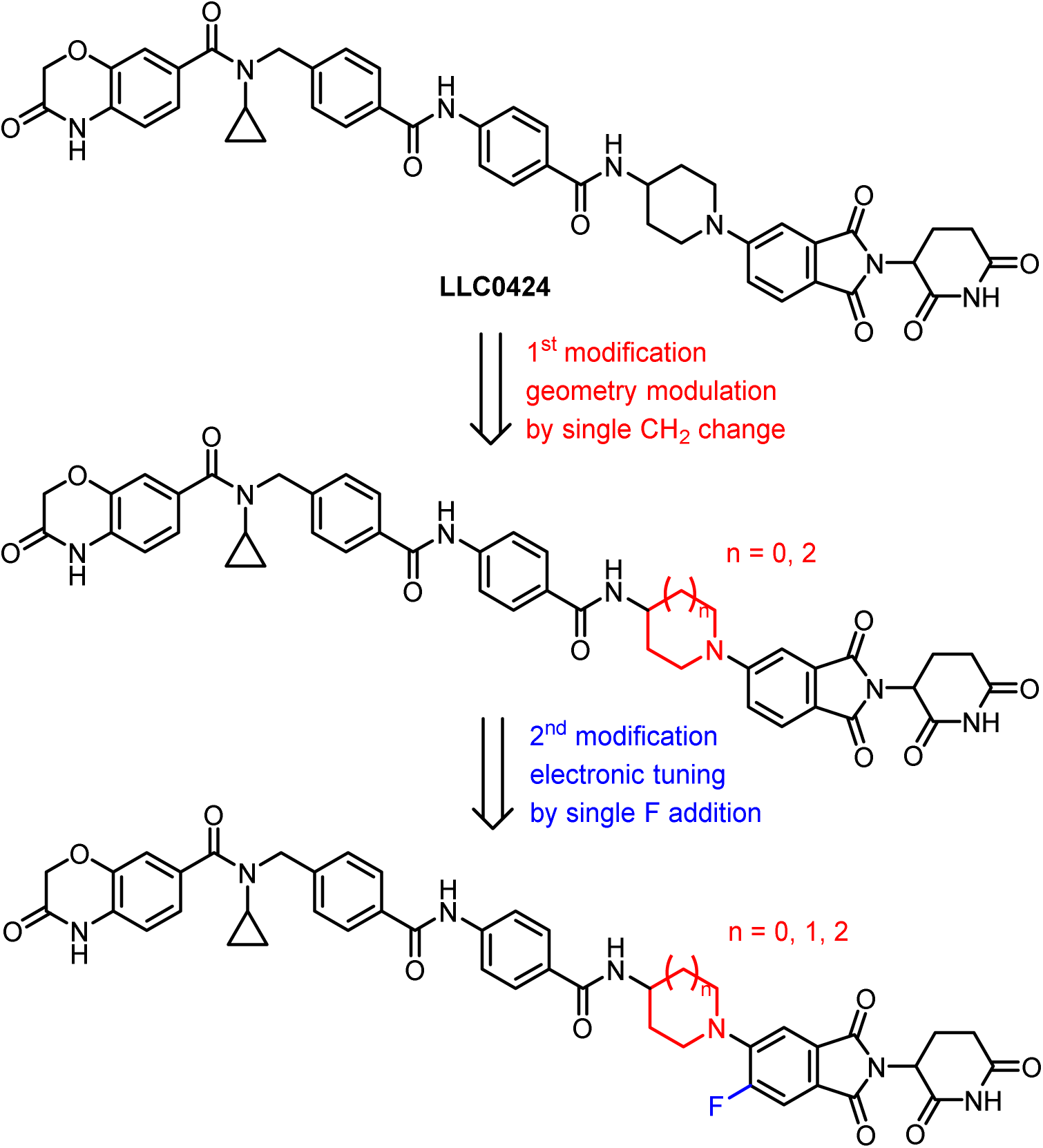
Molecular modification process on **LLC0424**.

Based on the modification strategy outlined above, we first designed compounds **8a** and **8b** by deleting or inserting a CH_2_ moiety in the piperidine linker. The degradation activity of these compounds was evaluated in prostate cancer VCaP cells, and the results revealed striking differences in their effects on target degradation. At a concentration of 2 μM, compound **8a** induced near-complete degradation of both NSD2 and GSPT1, with degradation levels of 97% and 98%, respectively. In contrast, compound **8b** potently degraded GSPT1 (98% degradation) while exhibiting markedly reduced activity against NSD2, with only 23% degradation observed. Subsequently, compounds **8c–8e** were synthesized by introducing a fluorine substituent at the 5′-position of the thalidomide moiety. The results demonstrated that fluorine substitution generally diminished the activity toward both NSD2 and GSPT1, as exemplified by compounds **8c** and **8d.** Notably, compound **8e** (**424-ND**), which differs from **LLC0424** by a single fluorine substitution, retained potent NSD2 degradation (97% at 2 μM) while showing no detectable degradation of GSPT1, indicating markedly improved selectivity. To further investigate the impact of stereochemistry on activity and selectivity, compound **8b** was resolved into its individual enantiomers. The *S*-enantiomer exerted minimal effects on both NSD2 and GSPT1, whereas the *R*-enantiomer (**424-GD**) exhibited pronounced GSPT1 degradation (reaching 99%) while maintaining significantly lower activity toward NSD2. Given the distinct degradation profiles of **424-ND** and **424-GD**, these two compounds were selected for further characterization.

**Table 1.**
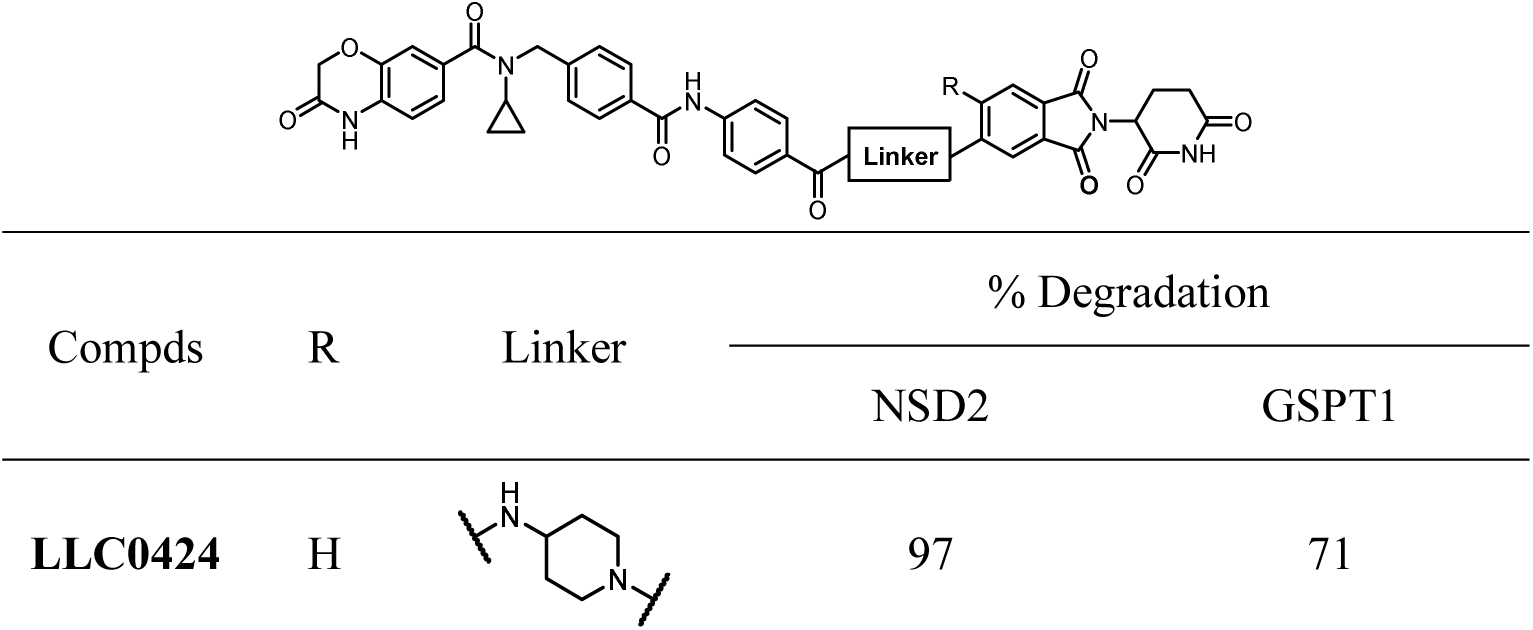
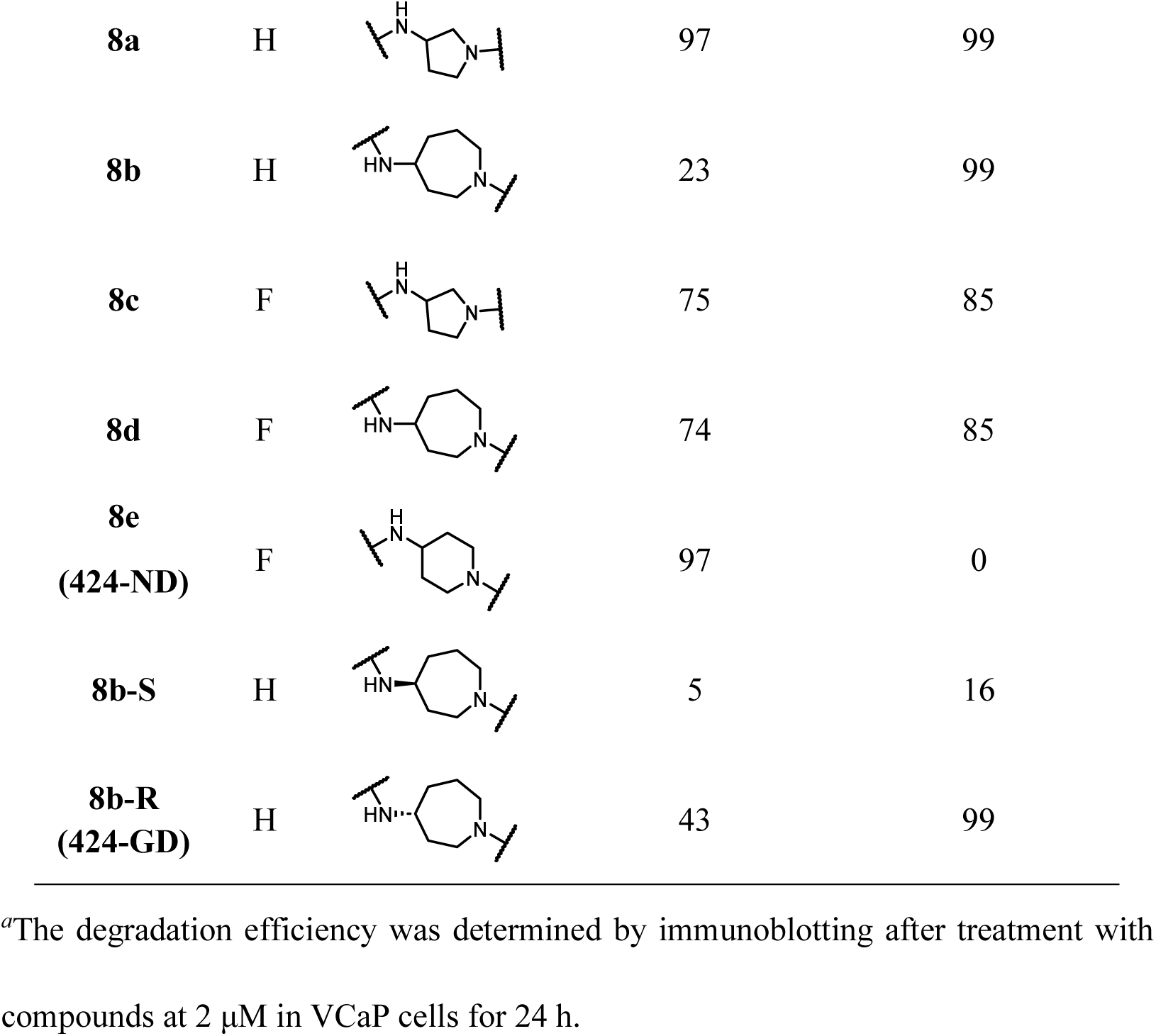
The Degradation Efficiency of Compounds 8a-e.*^a^*.

|  |  |  |  |  |
| --- | --- | --- | --- | --- |
| Compds | R | Linker | % Degradation |  |
|  |  |  | NSD2 | GSPT1 |
| **LLC0424** | H |  | 97 | 71 |
| **8a** | H |  | 97 | 99 |
| **8b** | H |  | 23 | 99 |
| **8c** | F |  | 75 | 85 |
| **8d** | F |  | 74 | 85 |
| **8e**<br>**(424-ND)** | F |  | 97 | 0 |
| **8b-S** | H |  | 5 | 16 |
| **8b-R**<br>**(424-GD)** | H |  | 43 | 99 |

<sup>a</sup>The degradation efficiency was determined by immunoblotting after treatment with compounds at 2 μM in VCaP cells for 24 h.

### 424-ND Selectively Induced NSD2 Degradation in a Dose-, CRBN– and Proteasome-Dependent Manner

To further characterize **424-ND**, we treated prostate cancer VCaP cells and multiple myeloma MM.1S cells with increasing concentrations of **424-ND** for 24 h. The results showed that **424-ND** dose-dependently degraded NSD2 by **424-ND** in both cell lines, whereas GSPT1 levels remained unchanged across the tested concentration range (Figure 3B,C). Tandem Mass Tag (TMT)-based quantitative proteomics analysis in VCaP cells further confirmed the selective downregulation of NSD2 upon **424-ND** treatment, with no widespread off-target protein degradation detected (Figure 3D). Both **424-ND-N1** (which lacks NSD2-binding capability) and **424-ND-N2** (which exhibits impaired CRBN engagement) failed to induce NSD2 degradation (Figure 3A,E). These findings indicate that engagement with both NSD2 and CRBN is essential for **424-ND**-mediated NSD2 degradation. Pretreatment with the NSD2 binder **UNC6934**, the CRBN ligand thalidomide, the NEDD8-activating enzyme inhibitor **MLN4924**, or the proteasome inhibitor bortezomib effectively abrogated **424-ND**-induced NSD2 degradation (Figure 3F,G,H,I). These results validate that **424-ND**-induced NSD2 degradation requires NSD2 engagement, CRBN recruitment, and an intact ubiquitin–proteasome system.

**Figure 3.**
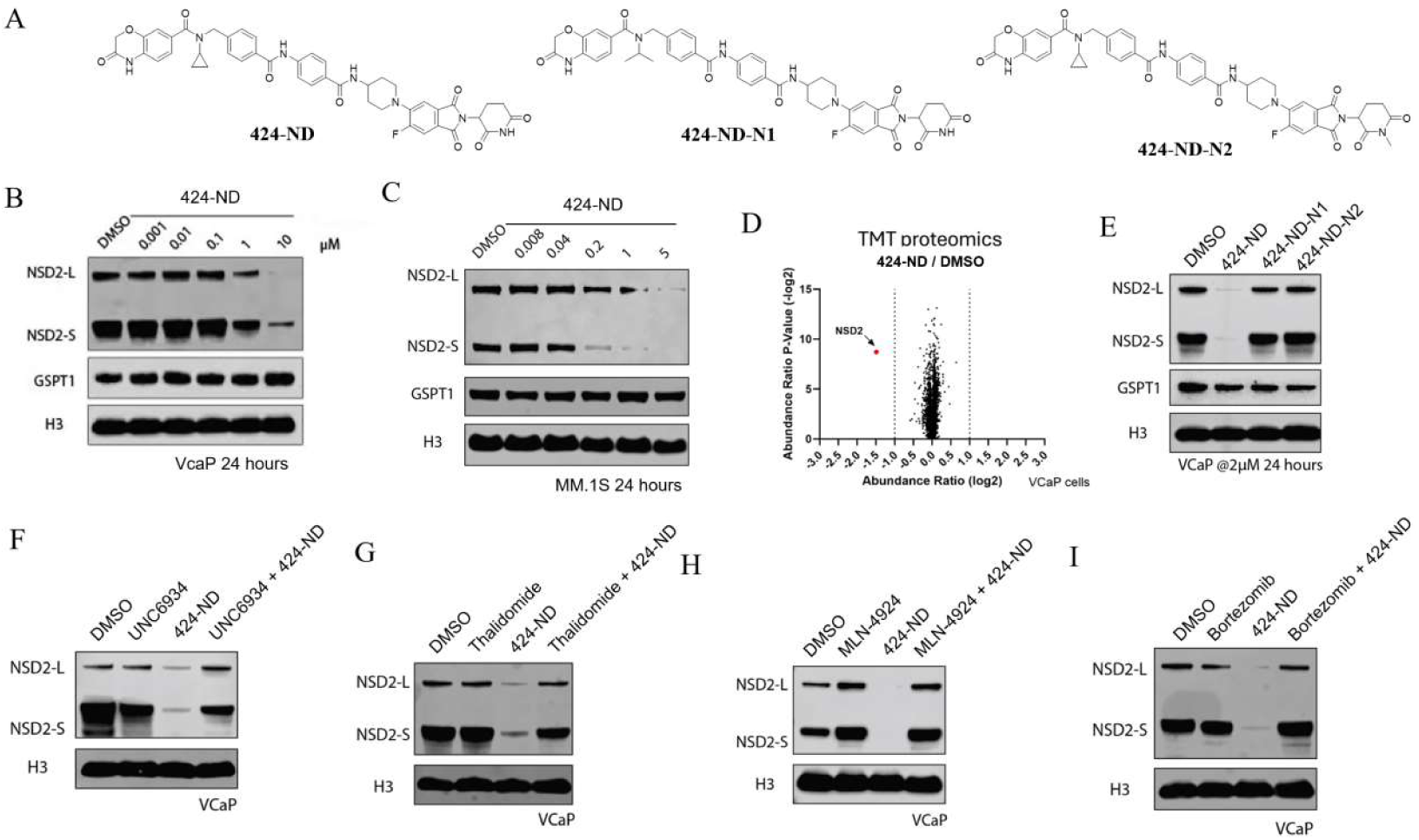
424-ND selectively induced NSD2 degradation in a dose-, CRBN– and proteasome-dependent manner. (A) Chemical structure of **424-ND**, **424-ND-N1** and **424-ND-N2**; Immunoblotting of NSD2 (long and short isoforms), GSPT1 and histone3 (H3) in VCaP cells (B) and MM.1S (C) treated with increasing concentrations of **424-ND** for 24 h; (D) Unbiased global proteomics analysis of **424-ND** in VCaP cells after 6 h treatment of DMSO or 2 μM **424-ND**; (E) Immunoblotting of NSD2 and H3 in VCaP cells treated with 2 µM **424-ND**, **424-ND-N1** and **424-ND-N2** for 24 h; Immunoblotting of NSD2 (long and short isoforms) and H3 in VCaP cells treated with 2 µM **424-ND** with or without NSD2 binder UNC6934 (F), CRBN ligand thalidomide (G), NEDD8-activating enzyme inhibitor MLN4924 (H) or proteasome inhibitor bortezomib (I).

### 424-GD Selectively Induced GSPT1 Degradation in a Dose-, CRBN– and Proteasome-Dependent Manner

As a selective GSPT1 degrader, **424-GD** induced robust, concentration-dependent degradation of GSPT1, while it exhibited significantly lower potency toward NSD2 (Figure 4B,C). Consistent with these cellular degradation profiles, TMT-based quantitative proteomic analysis in VCaP cells confirmed that treatment with **424-GD** led to pronounced downregulation of GSPT1, which in turn resulted in significant upregulation of ATF3 and GDF15, indicating activation of the integrated stress response (Figure 4D). Moreover, the NSD2-binding-deficient analogue **424-GD-N1** retained robust GSPT1 degradation activity, whereas the CRBN-binding-deficient analogue **424-GD-N2** failed to induce degradation of either GSPT1 or NSD2 (Figure 4A,E). Mechanistic studies demonstrated that pretreatment with the CRBN ligand thalidomide, the NEDD8-activating enzyme inhibitor **MLN4924**, or the proteasome inhibitor bortezomib markedly attenuated **424-GD**-induced GSPT1 degradation, whereas the NSD2 binder UNC6934 had no such effect (Figure 4F,G,H,I). These results indicate that **424-GD**-mediated GSPT1 degradation requires both CRBN engagement and proteasome activity.

**Figure 4.**
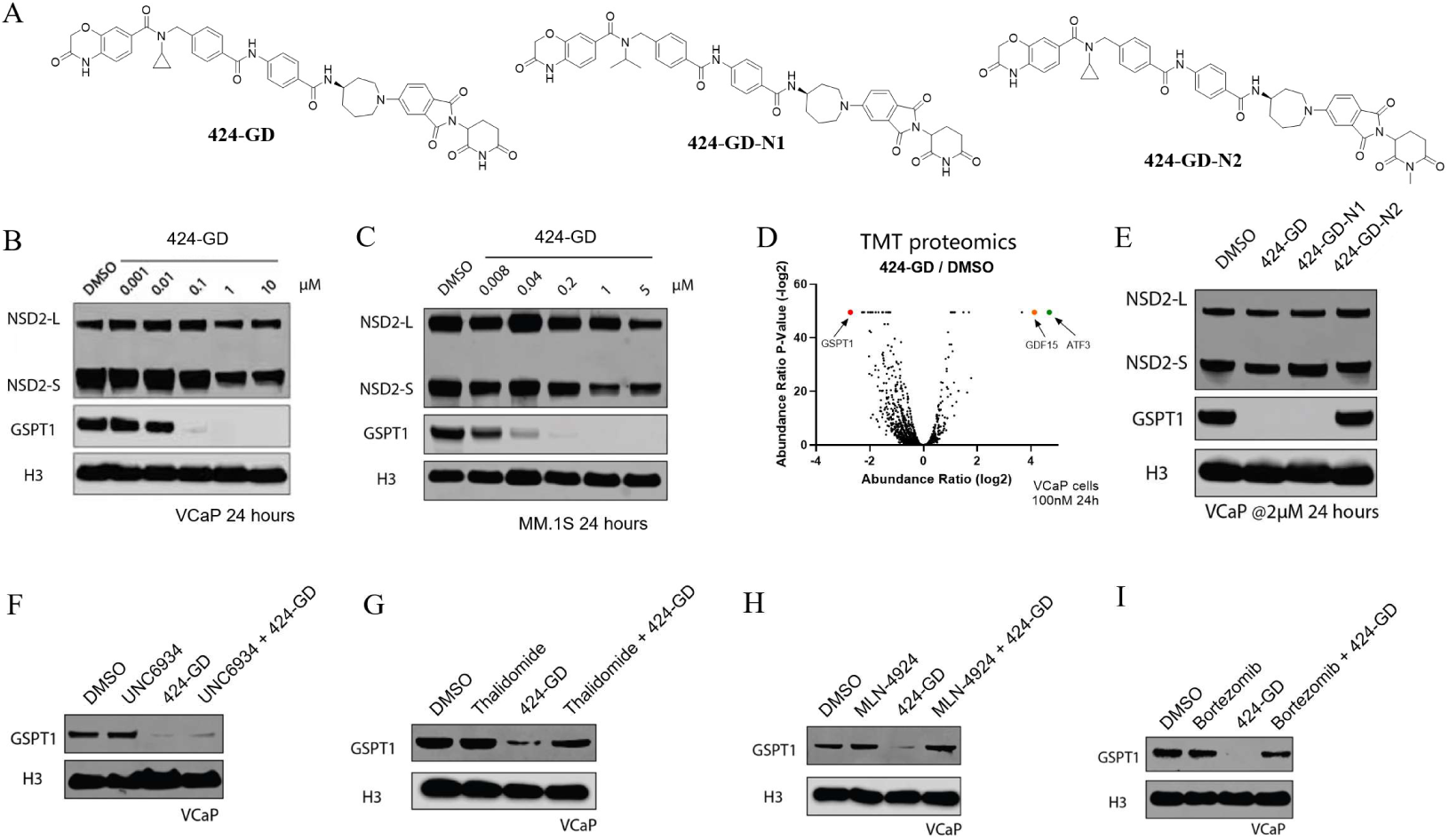
424-GD selectively induced GSPT1 degradation in a dose-, CRBN– and proteasome-dependent manner. (A) Chemical structure of **424-GD**, **424-GD-N1** and **424-GD-N2**; Immunoblotting of NSD2 (long and short isoforms), GSPT1 and histone3 (H3) in VCaP cells (B) and MM.1S (C) treated with increasing concentrations of **424-GD** for 24 h; (D) Unbiased global proteomics analysis of **424-GD** in VCaP cells after 24 h treatment of DMSO or 100 nM **424-GD**; (E) Immunoblotting of NSD2 and H3 in VCaP cells treated with 2 µM **424-GD**, **424-GD-N1** and **424-GD-N2** for 24 h; Immunoblotting of GSPT1 and H3 in VCaP cells treated with 2 µM **424-GD** with or without NSD2 binder UNC6934 (F), CRBN ligand thalidomide (G), NEDD8-activating enzyme inhibitor MLN4924 (H) or proteasome inhibitor bortezomib (I).

### Cell-based Phenotypic Evaluation of 424-ND and 424-GD

We further evaluated the cellular consequences associated with the selective degradation of NSD2 or GSPT1 by **424-ND** and **424-GD**, respectively. In VCaP cells, the selective GSPT1 degrader **424-GD** exhibited pronounced, dose-dependent suppression of cell viability, with an IC_50_ value of 0.13 μM, representing a substantial improvement in potency compared to the parent compound **LLC0424** (IC_50_ = 4.59 μM). In contrast, the selective NSD2 degrader **424-ND** exerted only a modest effect on cell viability (IC_50_ > 30 μM), which correlates with our previous findings that knockout of NSD2 caused mild cell growth inhibition^23^, suggesting that combination of NSD2 targeting with other proteins may be needed to achieve significant cytotoxic effects (Figure 5A). We next examined the impact of prolonged **424-ND** treatment on androgen receptor signaling pathways in VCaP cells. After 14 days of treatment, NSD2 protein levels remained markedly decreased, while GSPT1 abundance remained unchanged (Figure 5B). Consequently, expression of the AR downstream target genes including FKBP5 and KLK3 was substantially decreased, indicating functional suppression of the AR signaling following NSD2 degradation, which phenocopies NSD2 knockout in AR^+^ prostate cancer cells^23^. In contrast, the selective GSPT1 degrader **424-GD** expectedly induced robust upregulation of the integrated stress response markers ATF4 and ATF3 (Figure 5C)^19^. Consistently, **424-GD**, but not **424-ND**, markedly increased ATF4 and ATF3 levels concomitant with GSPT1 degradation (Figure S3), further supporting activation of the integrated stress response as a functional consequence of GSPT1 degradation. Collectively, although **424-ND** and **424-GD** are derived from the same degrader scaffold, their distinct degradation selectivity translates into markedly different cellular phenotypes and pathway-level outcomes.

**Figure 5.**
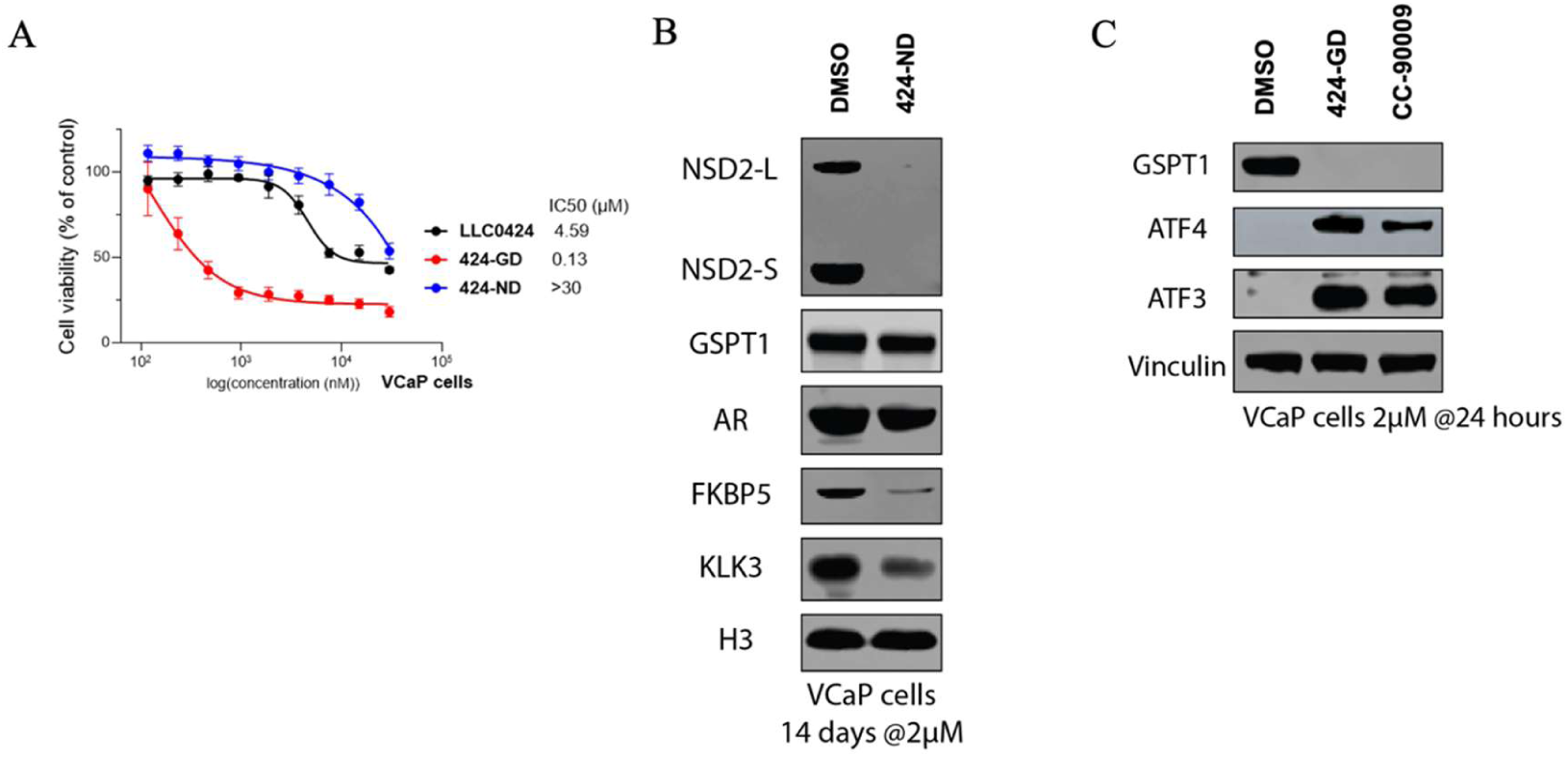
Distinct biological functions of **424-ND** and **424-GD**. (A) VCaP cells were treated with **LLC0424**, **424-GD** or **424-ND** at varying concentrations for 7 days and measured for cell viability by CellTiter-Glo assay. Data are reported as the mean of six independent experiments ± SD; (B) Immunoblot analysis of NSD2 (long and short isoforms), GSPT1, androgen receptor (AR), FKBP5, and KLK3 in VCaP cells treated with 2 μM **424-ND** or DMSO for 14 days. Histone H3 was used as a loading control; (C) Immunoblot analysis of GSPT1, ATF4 and ATF3 in VCaP cells treated with 2 μM of **424-GD** for 24 h. **CC-90009** (2 μM) was included as a reference compound.

### BLI Analysis of Ternary Complex Formation

To explore the distinct selectivity profiles arising from Single-atom modifications, biolayer interferometry (BLI) was employed to characterize compound-mediated modulation of ternary complex formation. Binary binding affinities between CRBN and the test compounds were first assessed. All compounds exhibited comparable apparent dissociation constants (*K*_d_) ranging from 120 to 160 nM, with consistent kinetic profiles across the tested concentration ranges—indicating that the structural modifications did not significantly alter intrinsic CRBN binding (Figure 6A,B,C). Markedly divergent binding behaviors were observed upon addition of GSPT1. Relative to the binary state, **LLC0424** and **424-ND** elicited only modest changes in CRBN binding responses, whereas **424-GD** induced a pronounced enhancement in binding signal accompanied by slower dissociation kinetics (Figure 6D,E,F). In line with this, cooperativity factor analysis (α = *K*_d_(binary)/*K*_d_(ternary)) revealed positive cooperativity for **424-GD** (α = 1.23), while **LLC0424** and **424-ND** yielded α values < 1. This cooperativity trend correlates with the superior GSPT1 degradation activity of **424-GD**. Conversely, a distinct binding pattern emerged in the presence of NSD2. Both **LLC0424** and **424-ND** displayed enhanced CRBN binding responses relative to the binary condition, with **424-ND** driving the most robust increase in binding stability (Figure 6G,H,I) and a cooperativity factor of 2.00—consistent with its elevated NSD2 degradation activity. In contrast, **424-GD** exhibited attenuated CRBN binding in the presence of NSD2, with an apparent *K*_d_ of 560 nM. Collectively, these BLI data demonstrate that subtle linker modifications profoundly impact ternary complex assembly, thereby differentially modulating functional degradation activity toward distinct target proteins.

**Figure 6.**
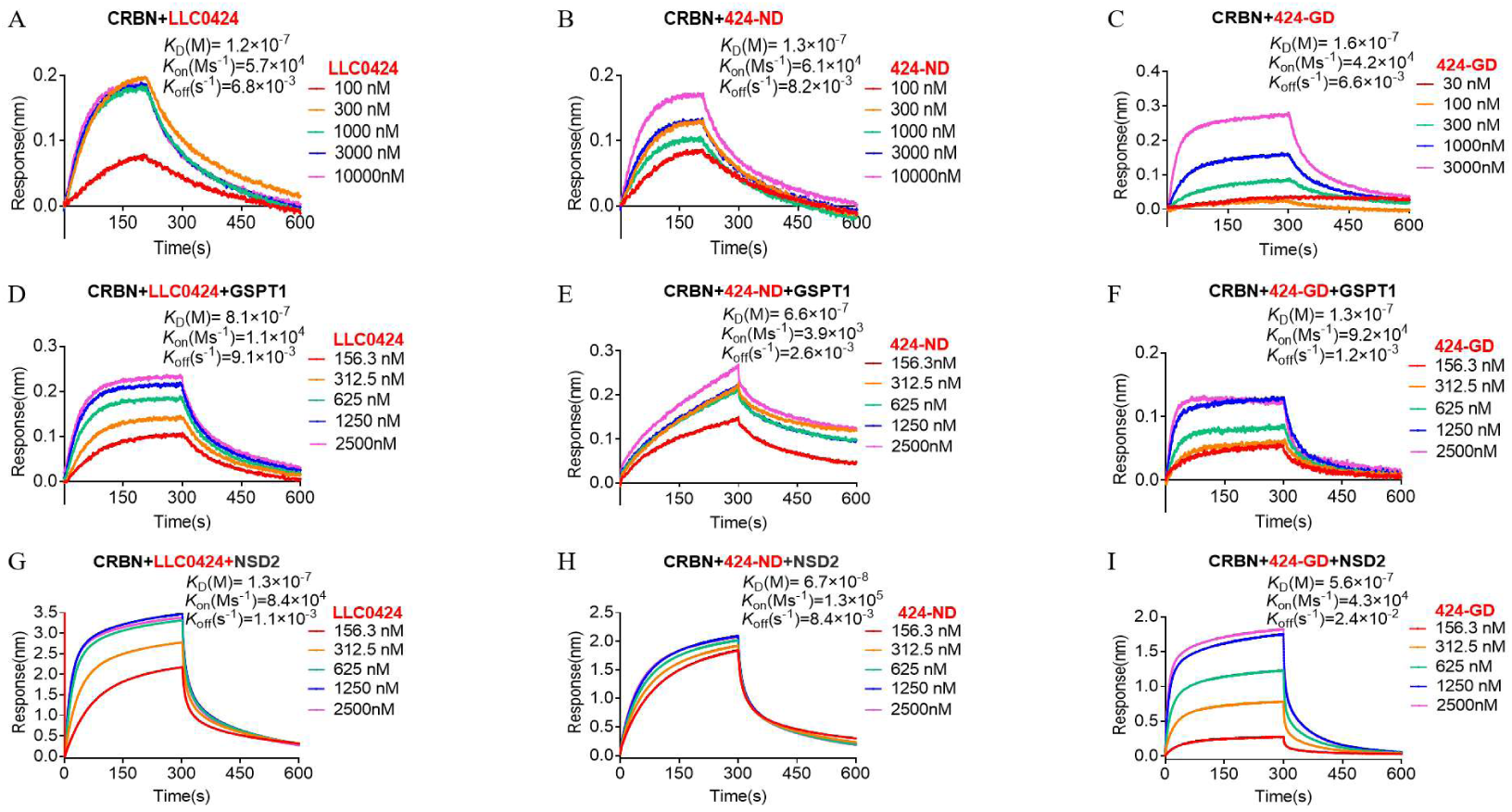
BLI analysis of CRBN–compound interactions in the presence of GSPT1 or NSD2. (A–C) BLI sensorgrams of CRBN binding to **LLC0424** (A), **424-ND** (B), and **424-GD** (C) under binary conditions. Biotinylated CRBN was immobilized on the sensor followed by incubation with different concentrations of compound. (D–F) BLI sensorgrams of CRBN binding to **LLC0424** (D), **424-ND** (E), and **424-GD** (F) in the presence of GSPT1. Biotinylated CRBN was immobilized on the sensor followed by incubation with designated concentrations of compound which was preincubated with GSPT1. (G–I) BLI sensorgrams of CRBN binding to **LLC0424** (G), **424-ND** (H), and **424-GD** (I) in the presence of NSD2. Biotinylated CRBN was immobilized on the sensor followed by incubation with designated concentrations of compound which was preincubated with NSD2.

### Molecular Dynamics Simulations Reveal Distinct Conformational Preferences of the Distal Moieties in Compound–CRBN Complexes

To further elucidate the molecular basis underlying the selectivity of these compounds, we modeled the compound-CRBN complexes based on previously reported crystal structures of IMiD-CRBN complexes (PDB ID: 5HXB), followed by long-timescale molecular dynamics simulations for structural equilibration. Interestingly, during the simulations, we observed notable differences in the dynamics of the solvent-exposed distal moieties of the compounds (Figure S2). To further characterize these conformational differences, we performed metadynamics simulations as described in the Methods section. Analysis of the converged two-dimensional free energy surfaces revealed that differences in linker structure significantly influenced the conformational distribution of the compounds, with each displaying distinct preferences (Figure 7). For **LLC0424**, seven local minima, designated M1-M7, were identified on the free energy surface. These minima mainly corresponded to two representative orientations of the **UNC6934** moiety: one in which the compound extended toward the solvent-exposed region between α1-helix β4-β5 loop of the CRBN surface, represented by M1-M4, and another in which the compound projected toward the solvent-exposed region close to β4 and β5, represented by M5-M7. In contrast, **424-ND** and **424-GD** preferentially sampled only one of these two orientations. Specifically, **424-ND** exhibited four major local minima, M1′-M4′, corresponding to the orientation represented by M1–M4 in **LLC0424**, whereas 424-GD displayed three major local minima, M1″-M3″, corresponding to the alternative orientation represented by M5-M7. These results suggest that differences in the conformational preferences of the solvent-exposed distal moieties may contribute to the distinct POI selectivity of **LLC0424**, **424-ND** and **424-GD,** providing a structural basis for the observed inversion of degradation selectivity.

**Figure 7.**
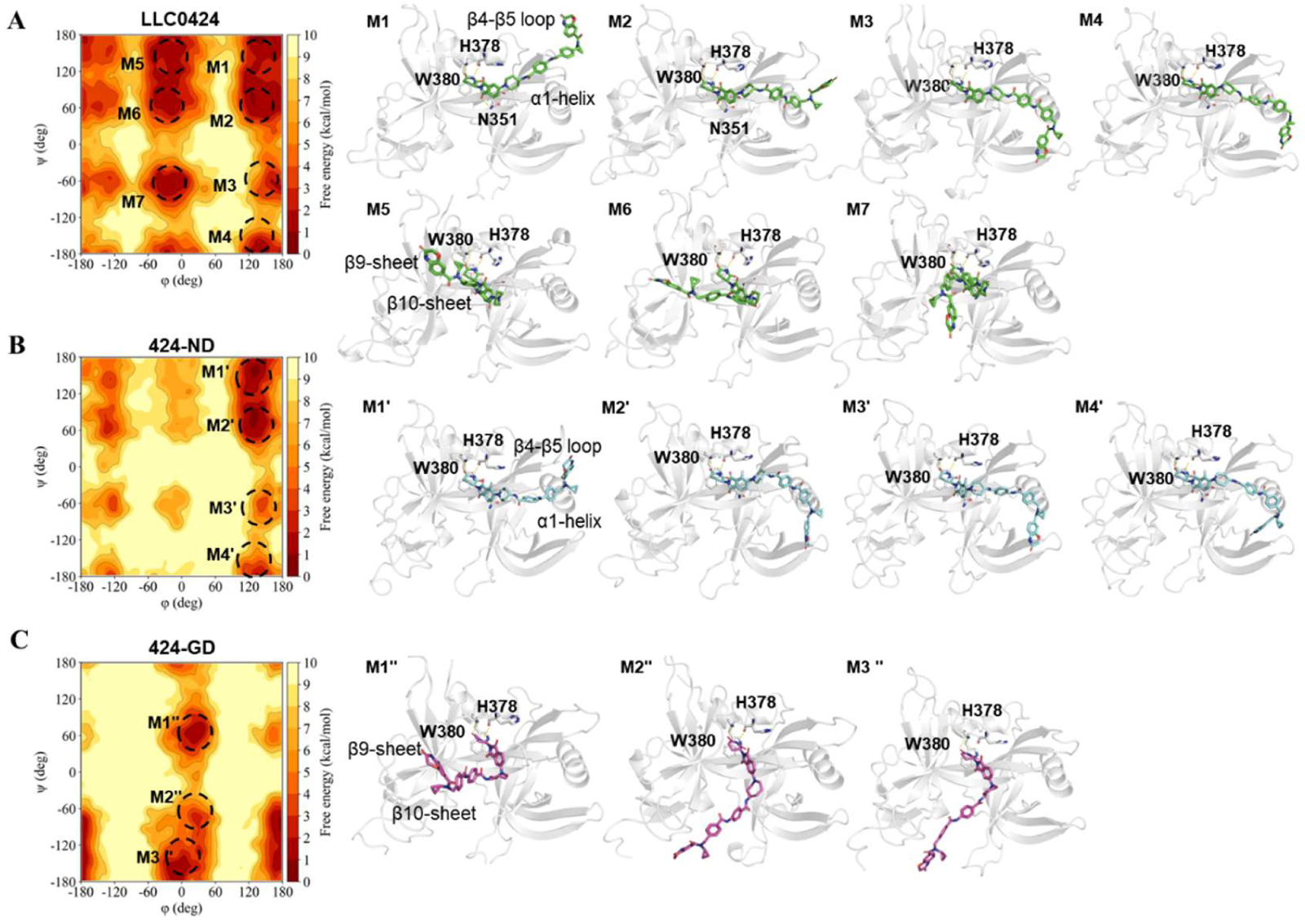
Conformational differences between **LLC0424**. (A), **424-ND** (B), and **424-GD** (C) in complex with CRBN from metadynamic simulations (using PDB: 5HXB as the model). The left panels show the two-dimensional free energy surfaces and the major local minima identified for each compound. The right panels show representative structures extracted from the corresponding local minima.

### Chemical Synthesis

The synthesis of compounds **8a–e** is outlined in Scheme 1. Briefly, a series of in-house Boc-protected thalidomide derivatives were first subjected to Boc deprotection, followed by amide coupling with 4-(4-((*N*-cyclopropyl-3-oxo-3,4-dihydro-2*H*-benzo[*b*][1,4]oxazine-7-carboxamido)methyl)benzamido)benzoic acid, yielding the target compounds **8a–e**.

**Scheme 1.**
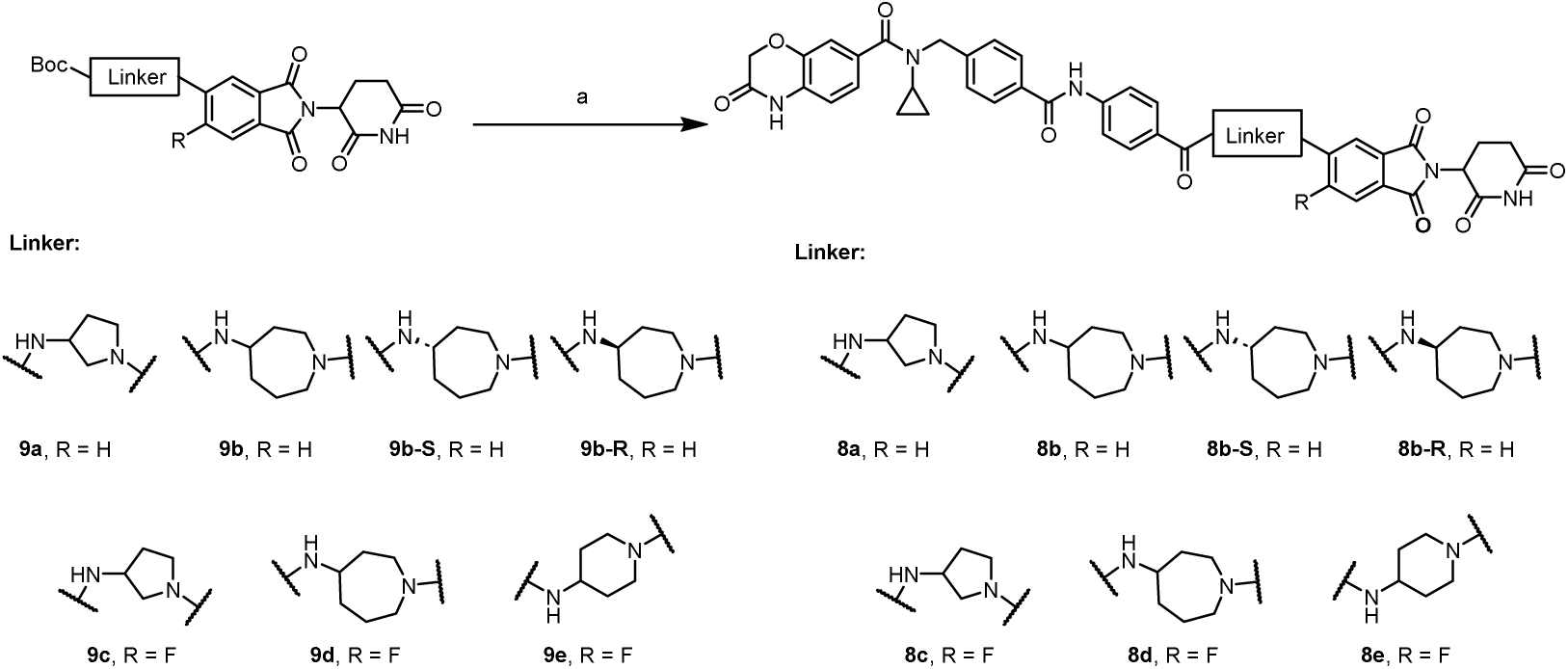
Synthesis of Compounds **8a-e***^a^*. *^a^* Reagents and conditions: (a) 1) trifluoroacetic acid (TFA), dichloromethane (CH_2_Cl_2_), room temperature (rt), 2 h; 2) 4-(4-((*N*-cyclopropyl-3-oxo-3,4-dihydro-2*H*-benzo[*b*][1,4]oxazine-7-carboxamido)methyl)benzamido)benzoic acid, *N*-[(dimethylamino)-3-oxo-1*H*-1,2,3-triazolo[4,5-*b*]pyridin-1-yl-methylene]-*N*-methylmethanaminium hexafluorophosphate (HATU), triethylamine (Et_3_N), *N, N*-dimethylformamide (DMF), rt, 2 h, 64-84% (two steps).

## CONCLUSION

Herein, single-atom modification of **LLC0424**, an NSD2 degrader with GSPT1 off-target activity, yields the NSD2-selective degrader **424-ND** *via* fluorine substitution and the GSPT1-selective degrader **424-GD** *via* methylene insertion. BLI, molecular dynamics and metadynamics simulations revealed that these functional differences arise from compound-specific ternary complex formation behaviors, which are tightly linked to the distinct conformational landscapes adopted by the degraders. While linker optimization is known to influence degrader activity, achieving dramatic selectivity inversion through such minimal structural changes remains highly unusual. These findings highlight the sensitivity of degrader selectivity to minimal local structural modifications and provide insights into the structural determinants governing selectivity in TPD.

From a functional perspective, as the biological roles of NSD2 remain incompletely defined, the development of selective NSD2 degraders holds significant value. Although several NSD2 degraders have been reported, proteome-wide selectivity remains uncharacterized for most. The degrader **424-ND** displays high selectivity across multiple cell lines, maintains selectivity under prolonged treatment, and modulates AR signaling consistent with NSD2 inhibition, supporting its utility as a high-quality chemical probe. Furthermore, the discovery of **424-GD** expands the chemical space for GSPT1 degrader development and provides new insights for the design of selective GSPT1-targeted degraders in drug discovery.

In summary, this work not only delivers highly selective degraders for NSD2 and GSPT1 but also highlights the remarkable sensitivity of degrader selectivity to minimal structural changes.

## EXPERIMENTAL SECTION

### General Chemistry Methods

The reagents and solvents used in chemical synthesis were obtained from commercial agents without further purification. All reactions were monitored by using thin-layer chromatography (TLC). All final compounds were purified by a column chromatography on silica gel (300-400 mesh). The NMR spectra were recorded on an Agilent DD2 500 MHz, or a Bruker Avance III HD 600 MHz NMR spectrometer in DMSO-*d*_6_. The spectra of high-resolution mass (HRMS) were monitored by Bruker MaXis 4G TOF Mass Spectrometer and an ESI source. The purities of all final compounds were identified by HPLC analysis with the Agilent 1200 system, and were proved to be >95%. HPLC condition: Triart C_18_ reversed-phase column, 5 μm, 4.6 mm × 250 mm, and flow rate 1.0 mL/min, starting with a 15 min-gradient from 0.1% TFA in water and acetonitrile 9:1 mixture to 0.1% TFA in acetonitrile, then ending with 0.1% TFA in acetonitrile for 5 min.

*N-cyclopropyl-N-(4-((4-((1-(2-(2,6-dioxopiperidin-3-yl)-1,3-dioxoisoindolin-5-yl)pyrrolidin-3-yl)carbamoyl)phenyl)carbamoyl)benzyl)-3-oxo-3,4-dihydro-2H-benzo[b][1,4]oxazine-7-carboxamide (**8a**). tert*-butyl (1-(2-(2,6-dioxopiperidin-3-yl)-1,3-dioxoisoindolin-5-yl)pyrrolidin-3-yl)carbamate (44.2 mg, 0.100 mmol) was dissolved in 2.0 mL of CH_2_Cl_2_. To above solution was added TFA (0.5 mL) by syringe. The mixture was stirred at room temperature for 2 h. After the reaction was complete, the solvent was removed by vacuum. The crude product was dissolved in a small amount of DMF (2.0 mL) and were added 4-(4-((*N*-cyclopropyl-3-oxo-3,4-dihydro-2*H*-benzo[*b*][1,4]oxazine-7-carboxamido)methyl)benzamido)benzoic acid (51.3 mg, 0.106 mmol), triethylamine (20.2 mg, 28 μL, 0.200 mmol), and HATU (45.6 mg, 0.120 mmol). The mixture was stirred at room temperature for 3 h. The resulting mixture was purified by column chromatography to afford the title compound as a yellow solid (62.3 mg, 77% yield). ^1^H NMR (500 MHz, DMSO-*d*_6_) δ 11.07 (s, 1H), 10.88 (s, 1H), 10.44 (s, 1H), 8.56 (d, *J* = 6.6 Hz, 1H), 7.96 (d, *J* = 8.2 Hz, 2H), 7.88 (s, 4H), 7.67 (d, *J* = 8.5 Hz, 1H), 7.46 (d, *J* = 7.9 Hz, 2H), 7.21 – 7.17 (m, 1H), 7.15 (d, *J* = 1.7 Hz, 1H), 6.95 (d, *J* = 2.2 Hz, 1H), 6.93 (d, *J* = 8.0 Hz, 1H), 6.86 (dd, *J* = 8.7, 2.2 Hz, 1H), 5.06 (dd, *J* = 12.9, 5.4 Hz, 1H), 4.72 (s, 2H), 4.70 – 4.65 (m, 1H), 4.62 (s, 2H), 3.81 – 3.74 (m, 1H), 3.67 – 3.58 (m, 1H), 3.57 – 3.48 (m, 1H), 3.46 – 3.40 (m, 1H), 2.94 – 2.83 (m, 1H), 2.81 (s, 1H), 2.63 – 2.51 (m, 2H), 2.32 (tt, *J* = 13.6, 7.4 Hz, 1H), 2.15 (dq, *J* = 12.5, 6.4 Hz, 1H), 2.06 – 1.97 (m, 1H), 0.55 (d, *J* = 6.9 Hz, 2H), 0.48 (s, 2H). ^13^C NMR (126 MHz, DMSO-*d*_6_) δ 173.3, 170.6, 168.2, 167.7, 166.4, 166.1, 165.3, 152.4, 143.0, 142.8, 142.4, 134.5, 133.9, 132.3, 129.5, 128.9, 128.6, 128.5, 127.7, 125.4, 122.3, 119.7, 116.2, 115.9, 115.8, 115.6, 106.1, 67.2, 53.5, 49.7, 49.2, 46.8, 31.4, 31.2, 23.0, 22.7, 10.0. HRMS (*m/z*): [M + H]^+^ calcd for C_44_H_40_N_7_O_9_^+^ 810.2882, found 810.2888. HPLC purity: 97.22%.

### Compounds 8b-e,424-ND-N2 were prepared by a procedure similar to that used for compound 8a

*N-cyclopropyl-N-(4-((4-((1-(2-(2,6-dioxopiperidin-3-yl)-1,3-dioxoisoindolin-5-yl)azepan-4-yl)carbamoyl)phenyl)carbamoyl)benzyl)-3-oxo-3,4-dihydro-2H-benzo[b][1,4]oxazine-7-carboxamide (**8b**).* Yield, 84%. ^1^H NMR (500 MHz, DMSO-*d*_6_) δ 11.08 (s, 1H), 10.88 (s, 1H), 10.41 (s, 1H), 8.22 (d, *J* = 7.7 Hz, 1H), 7.95 (d, *J* = 8.2 Hz, 2H), 7.87 – 7.79 (m, 4H), 7.67 (d, *J* = 8.5 Hz, 1H), 7.46 (d, *J* = 7.9 Hz, 2H), 7.22 – 7.17 (m, 1H), 7.14 (dd, *J* = 4.0, 2.0 Hz, 2H), 7.05 (s, 1H), 6.93 (d, *J* = 8.1 Hz, 1H), 5.07 (dd, *J* = 12.9, 5.4 Hz, 1H), 4.72 (s, 2H), 4.62 (s, 2H), 3.99 – 3.88 (m, 1H), 3.82 – 3.74 (m, 1H), 3.69 (dd, *J* = 14.1, 5.0 Hz, 1H), 3.64 – 3.54 (m, 2H), 2.94 – 2.83 (m, 1H), 2.80 (s, 1H), 2.63 – 2.51 (m, 2H), 2.14 – 2.08 (m, 1H), 2.07 – 1.95 (m, 2H), 1.89 – 1.74 (m, 3H), 1.55 (q, *J* = 11.9 Hz, 1H), 0.59 – 0.51 (m, 2H), 0.48 (s, 2H). ^13^C NMR (126 MHz, DMSO-*d*_6_) δ 173.3, 170.6, 168.2, 167.6, 166.0, 165.3, 165.1, 153.9, 143.0, 142.8, 142.2, 134.8, 133.9, 132.3, 130.0, 128.9, 128.5, 128.4, 127.7, 125.7, 122.3, 119.7, 116.2, 115.8, 115.6, 115.5, 105.7, 67.2, 49.9, 49.7, 49.2 48., 46.1, 33.5, 32.8, 31.5, 24.0, 22.7, 10.0. HRMS (m/z): [M + H]^+^ calcd for C_46_H_44_N_7_O_9_^+^ 838.3195, found 838.3197. HPLC purity: 95.35%.

*N-cyclopropyl-N-(4-((4-((1-(2-(2,6-dioxopiperidin-3-yl)-6-fluoro-1,3-dioxoisoindolin-5-yl)pyrrolidin-3-yl)carbamoyl)phenyl)carbamoyl)benzyl)-3-oxo-3,4-dihydro-2H-benzo[b][1,4]oxazine-7-carboxamide (**8c**).* Yield, 91%. ^1^H NMR (500 MHz, DMSO-*d*_6_) δ 11.08 (s, 1H), 10.88 (s, 1H), 10.43 (s, 1H), 8.52 (d, *J* = 6.5 Hz, 1H), 7.96 (d, *J* = 7.8 Hz, 2H), 7.88 (s, 4H), 7.63 (d, *J* = 12.5 Hz, 1H), 7.46 (d, *J* = 7.9 Hz, 2H), 7.19 (d, *J* = 8.1 Hz, 1H), 7.15 (s, 1H), 7.09 (d, *J* = 7.6 Hz, 1H), 6.93 (d, *J* = 8.0 Hz, 1H), 5.12 – 5.03 (m, 1H), 4.72 (s, 2H), 4.62 (s, 2H), 4.60 – 4.49 (m, 1H), 3.92 (s, 1H), 3.74 (d, *J* = 8.6 Hz, 1H), 3.69 – 3.56 (m, 2H), 2.94 – 2.83 (m, 1H), 2.80 (s, 1H), 2.62 – 2.51 (m, 2H), 2.28 – 2.19 (m, 1H), 2.09 (dd, *J* = 11.9, 6.0 Hz, 1H), 2.05 – 1.98 (m, 1H), 0.57 – 0.52 (m, 2H), 0.48 (s, 2H). ^13^C NMR (126 MHz, DMSO-*d*_6_) δ 173.3, 170.5, 167.5, 166.9, 166.5, 166.1, 165.3, 153.6 (d, *J* = 247.5 Hz), 143.0, 142.4, 133.9, 132.3, 130.0, 129.5, 128.9, 128.6 (d, *J* = 11.7 Hz), 127.7, 122.3, 119.7, 117.8 (d, *J* = 8.6 Hz)., 115.7 (d, *J* = 25.6 Hz), 112.2 (d, *J* = 25.4 Hz), 109.8, 67.2, 55.4, 49.8, 49.4, 48.7, 31.4, 31.0, 22.7. HRMS (m/z): [M + H]^+^ calcd for C H N O ^+^ 828.2788, found 828.2781. HPLC purity: 98.43%.

*N-cyclopropyl-N-(4-((4-((1-(2-(2,6-dioxopiperidin-3-yl)-6-fluoro-1,3-dioxoisoindolin-5-yl)azepan-4-yl)carbamoyl)phenyl)carbamoyl)benzyl)-3-oxo-3,4-dihydro-2H-benzo[b][1,4]oxazine-7-carboxamide (**8d**).* Yield, 73%. ^1^H NMR (500 MHz, DMSO-*d*_6_) δ 11.09 (s, 1H), 10.88 (s, 1H), 10.41 (s, 1H), 8.25 (d, *J* = 7.8 Hz, 1H), 7.95 (d, *J* = 7.9 Hz, 2H), 7.85 (s, 4H), 7.66 (d, *J* = 13.0 Hz, 1H), 7.46 (d, *J* = 7.9 Hz, 2H), 7.30 (d, *J* = 7.6 Hz, 1H), 7.19 (d, *J* = 8.0 Hz, 1H), 7.15 (s, 1H), 6.93 (d, *J* = 8.0 Hz, 1H), 5.08 (dd, *J* = 12.8, 5.5 Hz, 1H), 4.72 (s, 2H), 4.62 (s, 2H), 4.01 (t, *J* = 7.6 Hz, 1H), 3.65 (d, *J* = 13.7 Hz, 2H), 3.52 (t, *J* = 12.8 Hz, 2H), 2.89 (ddd, *J* = 16.7, 13.6, 5.4 Hz, 1H), 2.80 (s, 1H), 2.63 – 2.51 (m, 2H), 2.24 – 1.78 (m, 6H), 1.64 (d, *J* = 11.5 Hz, 1H), 0.55 (d, *J* = 6.9 Hz, 2H), 0.48 (s, 2H). ^13^C NMR (126 MHz, DMSO-*d*_6_) δ 173.3, 171.1, 170.5, 167.4, 166.8, 166.0, 165.3, 165.2, 155.9, 153.9, 144.8, 144.8, 143.0, 142.8, 142.2, 133.9, 132.3, 130.0, 129.8, 128.9, 128.5, 128.4, 127.7, 122.3, 119.7, 119.3, 119.2, 115.8, 115.6, 113.0, 112.7, 111.1, 67.2, 51.8, 50.0, 49.4, 48.5, 48.4, 34.7, 33.2, 31.4, 24.7, 22.6, 10.0. HRMS (m/z): [M + H]^+^ calcd for C_46_H_43_N_7_O_9_F^+^ 856.3101, found 856.3077. HPLC purity: 99.94%.

*N-cyclopropyl-N-(4-((4-((1-(2-(2,6-dioxopiperidin-3-yl)-6-fluoro-1,3-dioxoisoindolin-5-yl)piperidin-4-yl)carbamoyl)phenyl)carbamoyl)benzyl)-3-oxo-3,4-dihydro-2H-benzo[b][1,4]oxazine-7-carboxamide (**8e**).* Yield, 76%. ^1^H NMR (500 MHz, DMSO-*d*_6_) δ 11.11 (s, 1H), 10.88 (s, 1H), 10.43 (s, 1H), 8.25 (d, *J* = 7.8 Hz, 1H), 7.96 (d, *J* = 8.2 Hz, 2H), 7.88 (s, 4H), 7.74 (d, *J* = 11.4 Hz, 1H), 7.50 (d, *J* = 7.5 Hz, 1H), 7.46 (d, *J* = 7.9 Hz, 2H), 7.19 (dd, *J* = 8.1, 1.8 Hz, 1H), 7.15 (d, *J* = 1.8 Hz, 1H), 6.93 (d, *J* = 8.1 Hz, 1H), 5.12 (dd, *J* = 12.8, 5.5 Hz, 1H), 4.72 (s, 2H), 4.62 (s, 2H), 4.09 – 4.00 (m, 1H), 3.67 (d, *J* = 11.8 Hz, 2H), 3.06 (t, *J* = 12.0 Hz, 2H), 2.95 – 2.84 (m, 1H), 2.81 (s, 1H), 2.64 – 2.52 (m, 2H), 2.09 – 1.99 (m, 1H), 1.98 – 1.91 (m, 2H), 1.79 – 1.70 (m, 2H), 0.55 (d, *J* = 6.9 Hz, 2H), 0.48 (s, 2H). ^13^C NMR (126 MHz, DMSO-*d*_6_) δ 173.2, 170.4, 167.2, 166.7, 166.7, 166.1, 165.6, 165.3, 158.8, 156.8, 146.0, 146.0, 143.0, 142.8, 142.3, 133.9, 132.3, 129.9, 129.3, 129.2, 128.9, 128.5, 128.5, 127.7, 123.6, 123.5, 122.4, 119.8, 115.8, 115.6, 114.4, 112.6, 112.4, 67.2, 49.6, 49.5, 48.1, 46.7, 31.7, 31.4, 23.0, 22.6, 10.0. HRMS (m/z): [M + H]^+^ calcd for C H N O ^+^ 842.2944, found 842.2948. HPLC purity: 95.10%.

*N-cyclopropyl-N-(4-((4-(((4S)-1-(2-(2,6-dioxopiperidin-3-yl)-1,3-dioxoisoindolin-5-yl)azepan-4-yl)carbamoyl)phenyl)carbamoyl)benzyl)-3-oxo-3,4-dihydro-2H-benzo[b][1,4]oxazine-7-carboxamide (**8b-S**)*. Yield, 66%. ^1^H NMR (500 MHz, DMSO-*d*_6_) δ 11.07 (s, 1H), 10.88 (s, 1H), 10.41 (s, 1H), 8.21 (d, *J* = 7.8 Hz, 1H), 7.95 (d, *J* = 8.2 Hz, 2H), 7.88 – 7.79 (m, 4H), 7.66 (d, *J* = 8.5 Hz, 1H), 7.45 (d, *J* = 7.9 Hz, 2H), 7.21 – 7.16 (m, 1H), 7.14 (dd, *J* = 4.8, 2.0 Hz, 2H), 7.05 (dd, *J* = 8.7, 2.4 Hz, 1H), 6.93 (d, *J* = 8.1 Hz, 1H), 5.07 (dd, *J* = 12.9, 5.4 Hz, 1H), 4.71 (s, 2H), 4.62 (s, 2H), 3.94 (dd, *J* = 14.2, 7.7 Hz, 1H), 3.82 – 3.73 (m, 1H), 3.72 – 3.64 (m, 1H), 3.64 – 3.54 (m, 2H), 2.94 – 2.83 (m, 1H), 2.80 (s, 1H), 2.63 – 2.51 (m, 2H), 2.15 – 2.06 (m, 1H), 2.05 – 1.94 (m, 2H), 1.88 – 1.81 (m, 3H), 1.59 – 1.49 (m, 1H), 0.55 (d, *J* = 6.8 Hz, 2H), 0.47 (s, 2H). ^13^C NMR (126 MHz, DMSO-*d*_6_) δ 173.3, 171.1, 170.7, 168.2, 167.6, 166.0, 165.3, 165.1, 153.9, 143.0, 142.8, 142.2, 134.8, 133.9, 132.3, 130.0, 128.9, 128.5, 128.4, 127.7, 125.7, 122.3, 119.7, 116.2, 115.8, 115.6, 115.5, 105.7, 67.2, 50.7, 49.9, 49.7, 49.2, 48.1, 46.1, 33.5, 32.8, 31.5, 24.0, 22.7, 10.0. HRMS (m/z): calcd for C H N O ^+^ [M + H]^+^ 838.3195; found 838.3197. HPLC purity: 98.04%.

*N-cyclopropyl-N-(4-((4-(((4R)-1-(2-(2,6-dioxopiperidin-3-yl)-1,3-dioxoisoindolin-5-yl)azepan-4-yl)carbamoyl)phenyl)carbamoyl)benzyl)-3-oxo-3,4-dihydro-2H-benzo[b][1,4]oxazine-7-carboxamide (**8b-R**)*. Yield, 70%. ^1^H NMR (500 MHz, DMSO-*d*_6_) δ 11.07 (s, 1H), 10.88 (s, 1H), 10.41 (s, 1H), 8.21 (d, *J* = 7.8 Hz, 1H), 7.95 (d, *J* = 8.2 Hz, 2H), 7.89 – 7.79 (m, 4H), 7.66 (d, *J* = 8.5 Hz, 1H), 7.45 (d, *J* = 7.9 Hz, 2H), 7.21 – 7.16 (m, 1H), 7.14 (dd, *J* = 4.2, 2.0 Hz, 2H), 7.06 (dd, *J* = 8.8, 2.4 Hz, 1H), 6.93 (d, *J* = 8.0 Hz, 1H), 5.06 (dd, *J* = 12.9, 5.4 Hz, 1H), 4.71 (s, 2H), 4.61 (s, 2H), 4.01 – 3.89 (m, 1H), 3.82 – 3.73 (m, 1H), 3.72 – 3.65 (m, 1H), 3.66 – 3.53 (m, 2H), 2.94 – 2.83 (m, 1H), 2.80 (s, 1H), 2.63 – 2.51 (m, 2H), 2.09 (dd, *J* = 10.3, 5.8 Hz, 1H), 2.04 – 1.93 (m, 2H), 1.90 – 1.78 (m, 3H), 1.54 (q, *J* = 11.8 Hz, 1H), 0.57 – 0.51 (m, 2H), 0.47 (s, 2H). ^13^C NMR (126 MHz, DMSO-*d*_6_) δ 173.3, 171.1, 170.7, 168.2, 167.6, 166.0, 165.3, 165.1, 153.9, 143.0, 142.8, 142.2, 134.8, 133.9, 132.3, 130.0, 128.9, 128.5, 128.4, 127.7, 125.7, 122.3, 119.7, 116.2, 115.8, 115.6, 115.5, 105.7, 67.2, 49.9, 49.7, 49.2, 48.1, 46.1, 33.5, 32.8, 31.5, 24.0, 22.7, 10.0. HRMS (m/z): calcd for C_46_H_44_N_7_O_9_^+^ [M + H]^+^ 838.3195; found 838.3181. HPLC purity: 98.24%.

*N-(4-((4-((1-(2-(2,6-dioxopiperidin-3-yl)-6-fluoro-1,3-dioxoisoindolin-5-yl)piperidin-4-yl)carbamoyl)phenyl)carbamoyl)benzyl)-N-isopropyl-3-oxo-3,4-dihydro-2H-benzo[b][1,4]oxazine-7-carboxamide (**424-ND-N2**).* Yield, 80%. ^1^H NMR (500 MHz, DMSO-*d*_6_) δ 10.88 (s, 1H), 10.43 (s, 1H), 8.25 (d, *J* = 7.7 Hz, 1H), 7.96 (d, *J* = 8.2 Hz, 2H), 7.87 (s, 4H), 7.73 (d, *J* = 11.3 Hz, 1H), 7.50 (d, *J* = 7.4 Hz, 1H), 7.46 (d, *J* = 7.9 Hz, 2H), 7.19 (dd, *J* = 8.0, 1.7 Hz, 1H), 7.15 (d, *J* = 1.8 Hz, 1H), 6.93 (d, *J* = 8.1 Hz, 1H), 5.18 (dd, *J* = 13.1, 5.4 Hz, 1H), 4.72 (s, 2H), 4.62 (s, 2H), 4.04 (dq, *J* = 11.0, 5.5 Hz, 1H), 3.67 (d, *J* = 11.8 Hz, 2H), 3.06 (t, *J* = 12.0 Hz, 2H), 3.02 (s, 3H), 2.95 (ddd, *J* = 17.0, 13.8, 5.4 Hz, 1H), 2.81 – 2.72 (m, 2H), 2.61 – 2.51 (m, 1H), 2.10 – 2.02 (m, 1H), 1.98 – 1.92 (m, 2H), 1.79 – 1.69 (m, 2H), 0.58 – 0.52 (m, 2H), 0.48 (s, 2H). ^13^C NMR (126 MHz, DMSO-*d*_6_) δ 172.2, 170.0, 167.2, 166.7 (d, *J* = 2.5 Hz), 166.0, 165.6, 165.3, 158. 8, 156.8, 146.0 (d, *J* = 8.8 Hz), δ142.9 (d, *J* = 15.8 Hz), 142.3, 133.9, 132.3, 129.9, 129.2, 128.9, 128.5 (d, *J* = 6.3 Hz), 127.7, 123.5 (d, *J* = 9.8 Hz), 122.4, 119.8, 115.7 (d, *J* = 26.0 Hz), 114.5, 112.5 (d, *J* = 25.0 Hz). 67.2, 50.1, 49.6, 46.7, 31.7, 31.6, 27.1, 21.7. HRMS (m/z): [M + H]^+^ calcd for C_46_H_43_N_7_O_9_F^+^ 856.3101, found 856.3108.

*N-cyclopropyl-N-(4-((4-((1-(2-(2,6-dioxopiperidin-3-yl)-1,3-dioxoisoindolin-5-yl)azepan-4-yl)carbamoyl)phenyl)carbamoyl)benzyl)-3-oxo-3,4-dihydro-2H-benzo[b][1,4]oxazine-7-carboxamide (**424-GD-N2**).* Yield, 57%. ^1^H NMR (500 MHz, DMSO-*d*_6_) δ 10.88 (s, 1H), 10.41 (s, 1H), 8.21 (d, *J* = 7.8 Hz, 1H), 7.95 (d, *J* = 8.0 Hz, 2H), 7.88 – 7.79 (m, 4H), 7.66 (d, *J* = 8.5 Hz, 1H), 7.45 (d, *J* = 7.9 Hz, 2H), 7.19 (d, *J* = 8.1 Hz, 1H), 7.16 – 7.12 (m, 2H), 7.06 (dd, *J* = 8.7, 2.3 Hz, 1H), 6.93 (d, *J* = 8.1 Hz, 1H), 5.13 (dd, *J* = 13.0, 5.4 Hz, 1H), 4.71 (s, 2H), 4.61 (s, 2H), 4.00 – 3.87 (m, 1H), 3.81 – 3.73 (m, 1H), 3.72 – 3.66 (m, 1H), 3.62 – 3.53 (m, 2H), 3.01 (s, 3H), 2.99 – 2.91 (m, 1H), 2.85 – 2.73 (m, 2H), 2.62 – 2.52 (m, 1H), 2.15 – 2.07 (m, 1H), 2.06 – 2.01 (m, 1H), 2.01 – 1.93 (m, 1H), 1.88 – 1.77 (m, 3H), 1.60 – 1.49 (m, 1H), 0.57 – 0.52 (m, 2H), 0.49 – 0.45 (m, 2H). ^13^C NMR (126 MHz, DMSO-*d*_6_) δ 172.3, 171.1, 170.4, 168.2, 167.5, 166.0, 165.3, 165.1, 153.9, 143.0, 142.8, 142.2, 134.8, 133.9, 132.3, 130.0, 128.9, 128.5, 128.4, 127.7, 125.7, 122.3, 119.7, 116.2, 115.8, 115.6, 115.5, 105.7, 67.2, 49.9, 49.7, 49.7, 46.1, 33.5, 32.8, 31.6, 27.1, 24.0, 21.9, 10.0. HRMS (ESI m/z): calcd for C_47_H_46_N_7_O_9_^+^ [M + H]^+^ 852.3352; found 852.3352. HPLC purity: 99.82%.

*N-cyclopropyl-N-(4-((4-((1-(2-(2,6-dioxopiperidin-3-yl)-1,3-dioxoisoindolin-5-yl)azepan-4-yl)carbamoyl)phenyl)carbamoyl)benzyl)-3-oxo-3,4-dihydro-2H-benzo[b][1,4]oxazine-7-carboxamide (**424-ND-N1**). ter*t-butyl (1-(2-(2,6-dioxopiperidin-3-yl)-6-fluoro-1,3-dioxoisoindolin-5-yl)piperidin-4-yl)carbamate (47.4 mg, 0.531 mmol) was dissolved in CH_2_Cl_2_ (2.0 mL). To above solution was added TFA (2.0 mL) by syringe. The mixture was stirred at room temperature for 2 h. After the reaction was complete, the solvent was removed by vacuum. The crude product was dissolved in a small amount of DMF (2.0 mL) and were added 4-(4-((N-cyclopropyl-3-oxo-3,4-dihydro-2H-benzo[b][1,4]oxazine-7-carboxamido)methyl)benzamido)benzoic acid (284 mg, 0.584 mmol), triethylamine (161 mg, 0.22 mL, 1.594 mmol), and HATU (242 mg, 0.638 mmol). The mixture was stirred at room temperature for 3 h. The resulting mixture was purified by column chromatography to afford the title compound as a yellow solid. Yield, 78%. ^1^H NMR (400 MHz, DMSO-*d*_6_) δ 11.13 (s, 1H), 10.88 (s, 1H), 10.42 (s, 1H), 8.27 (d, *J* = 7.7 Hz, 1H), 7.92 (d, *J* = 8.1 Hz, 2H), 7.87 (s, 4H), 7.74 (d, *J* = 11.4 Hz, 1H), 7.50 (d, *J* = 7.3 Hz, 1H), 7.47 (s, 2H), 7.00 (m, 3H), 5.12 (dd, *J* = 12.9, 5.4 Hz, 1H), 4.63 (s, 4H), 4.05 (s, 2H), 3.70 – 3.62 (m, 2H), 3.05 (t, *J* = 11.7 Hz, 2H), 2.96 – 2.82 (m, 1H), 2.59 (d, *J* = 17.4 Hz, 2H), 2.09 – 2.00 (m, 1H), 1.94 (d, *J* = 12.3 Hz, 2H), 1.74 (d, *J* = 11.8 Hz, 2H), 1.10 (d, *J* = 6.5 Hz, 6H). ^13^C NMR (126 MHz, DMSO-*d*_6_) δ 173.2, 170.6, 170.4, 167.2, 166.7, 166.1, 165.6, 165.2, 157.8 (d, *J* = 253.0 Hz), 146.0 (d, *J* = 8.8 Hz), 144.2, 143.5, 142.3, 133.4, 132.3, 129.8, 129.3, 128.6, 128.5, 128.2, 127.0, 123.5 (d, *J* = 9.7 Hz), 121.0, 120.7, 116.2, 114.7, 114.4, 114.4, 112.4 (d, *J* = 25.2 Hz), 67.2, 49.6, 49.6, 49.5, 46.7, 43.3, 31.7, 31.4, 22.6, 21.2. HRMS (m/z): [M + H]^+^ calcd for C_45_H_43_N_7_O_9_F^+^ 844.3101, found 844.3089. HPLC purity: 97.70%.

*N-cyclopropyl-N-(4-((4-((1-(2-(2,6-dioxopiperidin-3-yl)-1,3-dioxoisoindolin-5-yl)azepan-4-yl)carbamoyl)phenyl)carbamoyl)benzyl)-3-oxo-3,4-dihydro-2H-benzo[b][1,4]oxazine-7-carboxamide (**424-GD-N1**).* Yield, 64%. ^1^H NMR (500 MHz, DMSO-*d*_6_) δ 11.07 (s, 1H), 10.86 (s, 1H), 10.38 (s, 1H), 8.21 (d, *J* = 7.8 Hz, 1H), 7.91 (d, *J* = 8.1 Hz, 2H), 7.86 – 7.78 (m, 4H), 7.66 (d, *J* = 8.5 Hz, 1H), 7.45 (d, *J* = 7.4 Hz, 2H), 7.14 (d, *J* = 2.3 Hz, 1H), 7.08 – 7.01 (m, 3H), 6.98 – 6.94 (m, 1H), 5.06 (dd, *J* = 12.9, 5.4 Hz, 1H), 4.62 (s, 4H), 4.16 – 4.02 (m, 1H), 3.98 – 3.87 (m, 1H), 3.81 – 3.74 (m, 1H), 3.71 – 3.65 (m, 1H), 3.64 – 3.53 (m, 2H), 2.94 – 2.83 (m, 1H), 2.67 – 2.53 (m, 2H), 2.15 – 2.06 (m, 1H), 2.05 – 1.92 (m, 2H), 1.90 – 1.77 (m, 3H), 1.59 – 1.49 (m, 1H), 1.10 (d, *J* = 6.8 Hz, 6H). ^13^C NMR (126 MHz, DMSO-*d*_6_) δ 173.3, 170.7, 168.2, 167.6, 166.1, 165.2, 165.1, 153.9, 143.4, 142.2, 134.8, 133.4, 132.3, 129.9, 128.6, 128.4, 128.2, 127.0, 125.7, 121.0, 119.7, 116.2, 115.5, 114.7, 105.7, 67.2, 49.9, 49.7, 49.2, 46.1, 33.5, 32.8, 31.5, 24.0, 22.7, 21.2. HRMS (m/z): calcd for C H N O ^+^ [M + H]^+^ 840.3352; found 840.3341. HPLC purity: 98.50%.

### Cell Lines, Antibodies, and Compounds

VCaP cells and MM.1S were obtained from ATCC. Both cell lines were tested for mycoplasma contamination every two weeks and were genotyped every three months by the University of Michigan Sequencing Core.. VCaP was maintained in Gibco DMEM (ThermoFisher) + 10% FBS (cytiya). MM.1S was maintained in RPMI 1640 Medium (ATCC modification) (Thermo Fisher) + 10% FBS (cytiva). Sources of all antibodies and compounds are included in Supplementary Table 1.

### Western Blot

After treatment, the cells were lysed in RIPA buffer (ThermoFisher Scientific) with Halt protease and phosphatase inhibitor cocktail (ThermoFisher Scientific). Protein quantification was determined by Pierce^TM^ Bovine Serum Albumin Standard Pre-Diluted Set (ThermoFisher Scientific). Equal amount of protein was denatured with LDS sample buffer (Invitrogen) supplemented with Sample Reducing Agent (Invitrogen) and resolved in NuPAGE 4–12% Bis-Tris protein gel (ThermoFisher Scientific). After transferring on Nitrocellulose membranes ((ThermoFisher Scientific) and blocking in 5% milk in 1X TBST, the membranes were incubated with primary antibodies overnight followed by HRP-conjugated secondary antibody incubation. The membranes were imaged on an Odyssey Fc Imager (LiCOR Biosciences). In the figures, one representative loading control was displayed.

### Compound Screening and Validation of Mechanism of Action

After the compounds were synthesized or purchased, they were diluted in DMSO. VCaP or MM.1S cells were plated into 6-well plates and were treated with the indicated compound. Western blot was then performed as previously described to probe for the certain proteins.

### TMT Proteomics Assay

VCaP cells were plated on 10 cm tissue culture plates overnight before treating with DMSO (control) or 2 µM **424-ND** with 3 replicates for each treatment. After 8 h, whole cell lysates were collected using RIPA buffer (ThermoFisher Scientific) and quantified using Pierce™ Bovine Serum Albumin Standard Pre-Diluted Set (ThermoFisher Scientific). Cell lysates were proteolyzed and labelled with TMT 6-plex Isobaric Label Reagent (ThermoFisher Scientific) according to manufacturer’s protocol and subjected to 8 fractions of liquid chromatography-mass spectrometry (LC-MS)/MS analysis as described The results were plotted with graphpad prism. To explore protein level changes with high confidence, the results of (LC-MS)/MS analysis was filtered with the criteria of peptides number > 4 and unique peptides number >1.

### Cell Viability Assay

VCaP cells were plated into 96-well plates and incubated overnight. A two-fold serial dilution of the indicated compounds were prepared in the culture medium and added to the cells the next day, with 6 replicates for each concentration, with 30 µM as the highest concentration being. The plate was further incubated for 7 days. Finally, the CellTiter-Glo assay (Promega) was performed according to the manufacturer’s protocol to determine cell proliferation and an Infinite M1000Pro microplate reader was used to acquired. The luminescence signal intensity. The data was analyzed using GraphPad Prism software.

### Biolayer Interferometry Assay

Purified recombinant CRBN was first biotinylated using the Thermo EZ-link long-chain biotinylation reagent. The CRBN and biotinylation reagent were then mixed in a 1:2 molar ratio in phosphate-buffered saline (PBS) at room temperature for 30 min. Then, the BLI binding assays were performed in 96-well (Greiner 655209) microplates at room temperature while shaking (1000 rpm) using the Octet R8 system (Sartorius). PBS with 0.1% bovine serum albumin (BSA), 0.05% Tween-20, and with or without 0.1% DMSO was used as assay buffer. Biotinylated CRBN protein was tethered on Super Streptavidin (SSA) biosensors (Sartorius) by dipping sensors into 200 μL of protein solutions (50 μg/mL/well). Averagely, a saturation response curve of 6−8 nm was achieved in 10 min. The measurement processes were all under computer control. Data were generated and analyzed by Octet User software. For the initial step, biosensors were washed in an assay buffer for 150 s to form a stable baseline.

1. Binary KD. The biosensors labeled with biotinCRBN were exposed to the solution containing the compound at the designated concentrations for the association and were monitored for 180 s. The biosensors were then moved back into the assay buffer to disassociate for another 420 s.
2. Ternary KD.The biosensors labeled with biotinCRBN were exposed to the solution containing the compound and protein complex for the association and were monitored for 300 s (100 μL compound in assay buffer added to 100 μL of 5 μM GSPT1 or NSD2). The biosensors were then moved back into the assay buffer to disassociate for another 300 s.

### Molecular Modeling

Complex structures of **LLC0424**, **424-ND**, and **424-GD** in complex with CRBN were generated based on the crystal structure (PDB ID: 5HXB) using molecular docking via Glide (Schrödinger, LLC, New York, NY, 2024) with default settings. Molecular dynamics (MD) simulations and metadynamics simulations were performed using Desmond. The OPLS4 force field and SPC water model were applied. Each system was initially relaxed for 5 ns, followed by three independent 500 ns production MD simulations in the NPT ensemble at 300 K and 1 atm using default settings. For metadynamics simulations, the dihedral angles of the compounds (as illustrated in Figure S1) were selected as collective variables (CVs). The Gaussian bias height was set to 0.3 kcal/mol and deposited every 0.09 ps. Each metadynamics simulation was run for a total of 500 ns to ensure adequate sampling of conformational space.

## ASSOCIATED CONTENT

### Supporting Information

**Figure S1** Definition of collective variables; **Figure S2** – Evolution of molecular dihedral with simulation time for **LLC0424**, **424-ND**, and **424-GD** across three independent molecular dynamics simulations; **Figure S3** – Immunoblot analysis of GSPT1, ATF4 and ATF3 in VCaP cells treated with 2 μM of **424-ND** and **424-GD** for 24 h, Sources of antibodies (**Table S1**) and compounds (**Table S2**); ^1^H NMR and ^13^C NMR spectra and HPLC traces for all degraders.

**Table S3** Molecular formula strings (CSV).

**Table S4** TMT proteomics data for 424-ND in VCaP cells (xlsx).

**Table S5** TMT proteomics data for 424-GD in VCaP cells (xlsx).

**Supplementary Data S1** Representative structures from metadynamic simulations extracted from the major local minima: LLC0424-CRBN (M1–M7), 424-ND-CRBN (M1–M4), 424-GD-CRBN (M1–M3) (PDB).

## AUTHOR INFORMATION

### Corresponding Authors

* (Y.Z.);

* (Z.W.);

* (A.M.C.);

*; Tel: +86-21-5492 5100 (K.D.)

### Author Contributions

*^#^*W. S., Y. L., L. L., Y. L., Y. Z., and Y. C. contributed equally to this work.

### Notes

K.D. is an advisor for Kinoteck Therapeutics and NuLynx Therapeutics. A.M.C. co-founded and serves on the scientific advisory boards of Lynx Dx, Medsyn Pharma, NuLynx Therapeutics, and Esanik Therapeutics.

## Supporting information

Supporting Information: Figures S1 – S3, Tables S1 – S2, NMR and HPLC traces

Supplemental Data S1: M1_LLC0424

Supplemental Data S1: M2_LLC0424

Supplemental Data S1: M3_LLC0424

Supplemental Data S1: M4_LLC0424

Supplemental Data S1: M5_LLC0424

Supplemental Data S1: M6_LLC0424

Supplemental Data S1: M7_LLC0424

## ACKNOWLEDGMENTS

We acknowledge the financial support from the National Natural Science Foundation of China (82530108 & 22521104), the National Key R&D Program of China (2023YFE0119000, 2023YFF1205104, 2023YFC2506402), the Strategic Priority Research Program of the Chinese Academy of Sciences (XDB1060000), the State Key Laboratory of Chemical Biology.

## ABBREVIATIONS USED

BLI: biolayer Interferometry
Boc: *t*-butyloxy carbonyl
CH_2_Cl_2_: dichloromethane
CRBN: cereblon
DMF: N,N-dimethylformamide
DMSO: dimethylsulfoxide
eRF1: eukaryotic release factor 1
eRF3a: eukaryotic peptide chain release factor 3a
Et_3_N: triethylamine
FKBP5: FK506 binding protein 5
GSPT1: G1 to S Phase Transition 1
GTP: guanosine triphosphate
H3K36me2: dimethylation of histone 3 lysine36
HATU: N-[(dimethylamino)-1H-1,2,3-triazolo[4,5-b]pyridin-1-ylmethylene]-N-methylmethanaminium hexafluorophosphate N-oxide
HPLC: high performance liquid chromatography
HRMS: high-resolution mass
IKZF1: Ikaros family zinc finger1
IKZF3: Ikaros family zinc finger3
KLK3: kallikrein related peptidase 3
NEDD8: neural precursor cell expressed, developmentally down-regulated 8
NMR: nuclear magnetic resonance
NSD2: Nuclear receptor-binding SET domain-containing 2
PROTAC: proteolysis targeting chimera
rt: room temperature
TFA: trifluoroacetic acid
TPD: targeted protein degradation

---

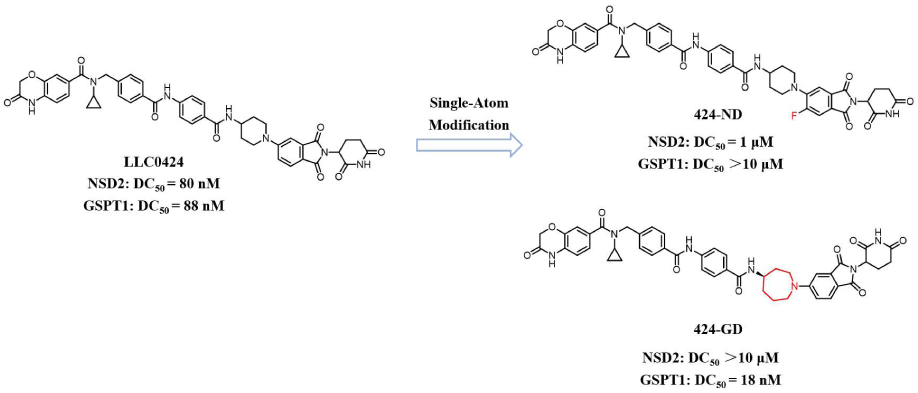
Table of Contents Graphic.

