## Supporting Information: Figures S1 – S3, Tables S1 – S2, NMR and HPLC traces for "Single-Atom Modification Enables Reversal of Degradation Selectivity between NSD2 and GSPT1"

### Table of Contents

**1.** Definition of collective variables for **LLC0424**, **424-ND**, and **424-GD**………….. S4

**2.** Evolution of molecular dihedral with simulation time for (A) **LLC0424**, (B) **424-ND**, and (C) **424-GD** across three independent molecular dynamics simulations….. S5

**3.** Immunoblot analysis of GSPT1, ATF4 and ATF3 in VCaP cells treated with 2 μM of **424-ND** and **424-GD** for 24 h………………………………………………………. S6

**4.** Sources of antibodies and compounds…………………………………………… S7

**5.** $^{1}H$ and $^{13}C$ NMR spectra for final compounds................................................ S8-18

**6.** HPLC spectra of final compounds................................................................ S19-23

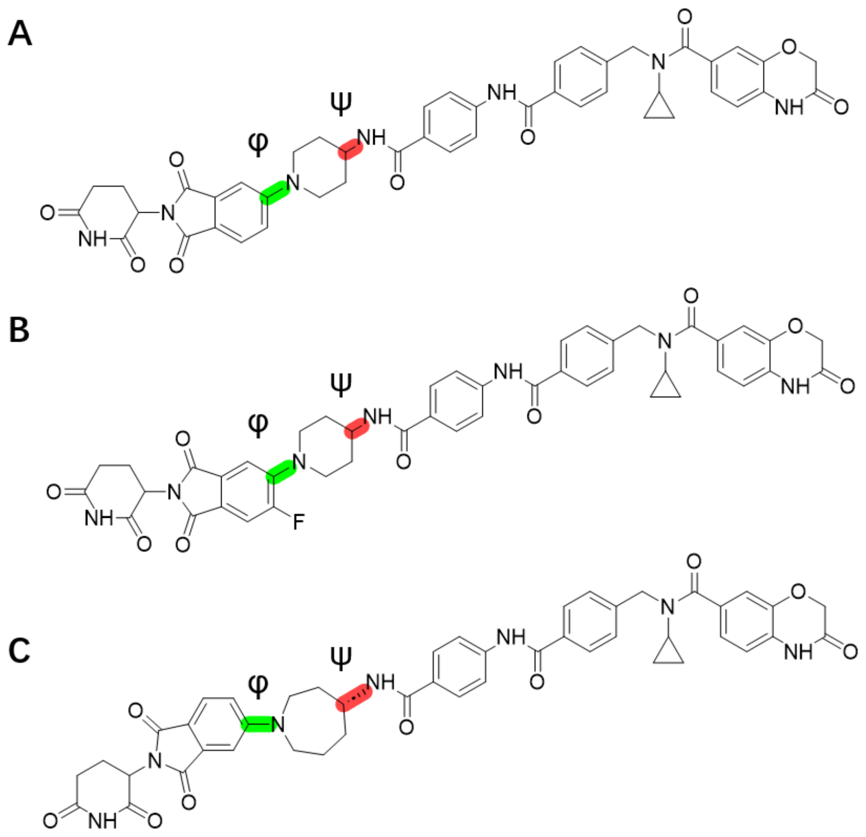


Figure S1. Definition of collective variables for (A) **LLC0424**, (B) **424-ND**, and (C) **424-GD**.

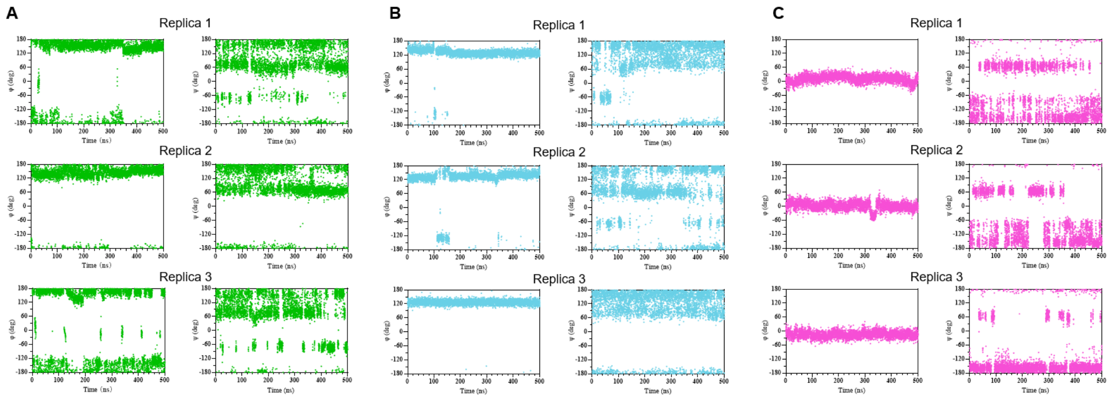


Figure S2. Evolution of molecular dihedral with simulation time for (A) LLC0424, (B) 424-ND, and (C) 424-GD across three independent molecular dynamics simulations.

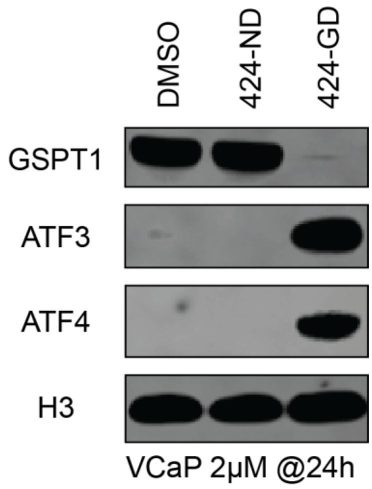


Figure S3. Immunoblot analysis of GSPT1, ATF4 and ATF3 in VCaP cells treated with 2 μM of **424-ND** and **424-GD** for 24 h.

**Table S1.** Sources of all antibodies.

| Primary antibodies used | | | | |
|---|---|---|---|---|
| **Antigen** | **Vendor** | **Catalog number** | **Application** | **Note** |
| NSD2 | Abcam | ab75359 | Western Blot | 1:1000 |
| eRF3 | Cell Signaling Technology | 14980 | Western Blot | 1:1000 |
| H3 | Cell Signaling Technology | 3638 | Western Blot | 1:2000 |
| AR | Abcam | ab108341 | Western Blot | 1:1000 |
| FKBP5 | Cell Signaling Technology | 12210 | Western Blot | 1:1000 |
| KLK3 | Cell Signaling Technology | 5365 | Western Blot | 1:1000 |
| ATF3 | Cell Signaling Technology | 33593 | Western Blot | 1:1000 |
| ATF4 | Cell Signaling Technology | 11815 | Western Blot | 1:1000 |
| **Secondary antibodies used** | | | | |
| **Product** | **Vendor** | **Catalog number** | **Application** | **Note** |
| Goat Anti-Rabbit IgG (H + L)-HRP Conjugate | Bio-Rad | 1706515 | Western Blot | 1:20000 |
| Goat Anti-Mouse IgG (H + L)-HRP Conjugate | Bio-Rad | 1706516 | Western Blot | 1:20000 |

**Table S2.** Sources of compounds.

| Name | Vendor | Catalog number |
|---|---|---|
| Thalidomide | Selleck Chemicals | S1193 |
| CC-90009 | MedChemExpress | HY-130800 |
| Bortezomib | Selleck Chemicals | S1013 |
| MLN4924 | Selleck Chemicals | S7109 |
| UNC6934 | Sigma-Aldrich | SML2813-5MG |

#### $^{1}H$ and $^{13}C$ NMR Spectra for final compounds

$^{1}H$ NMR and $^{13}C$ NMR spectra for **8a**

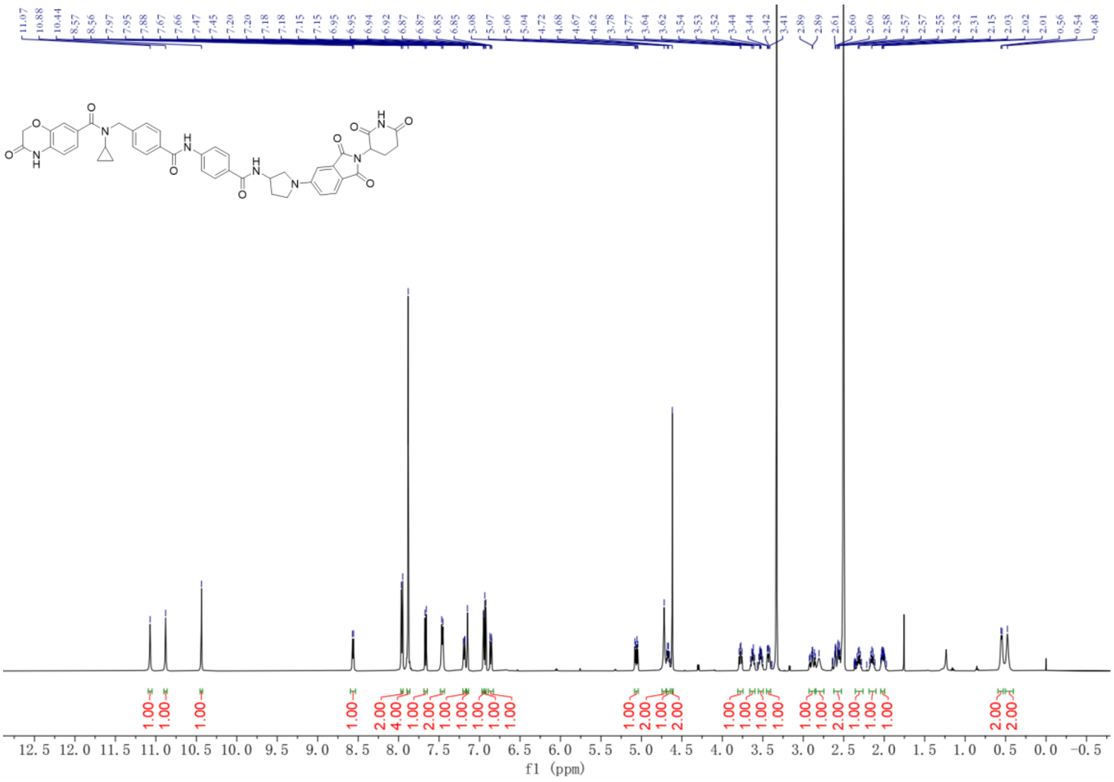


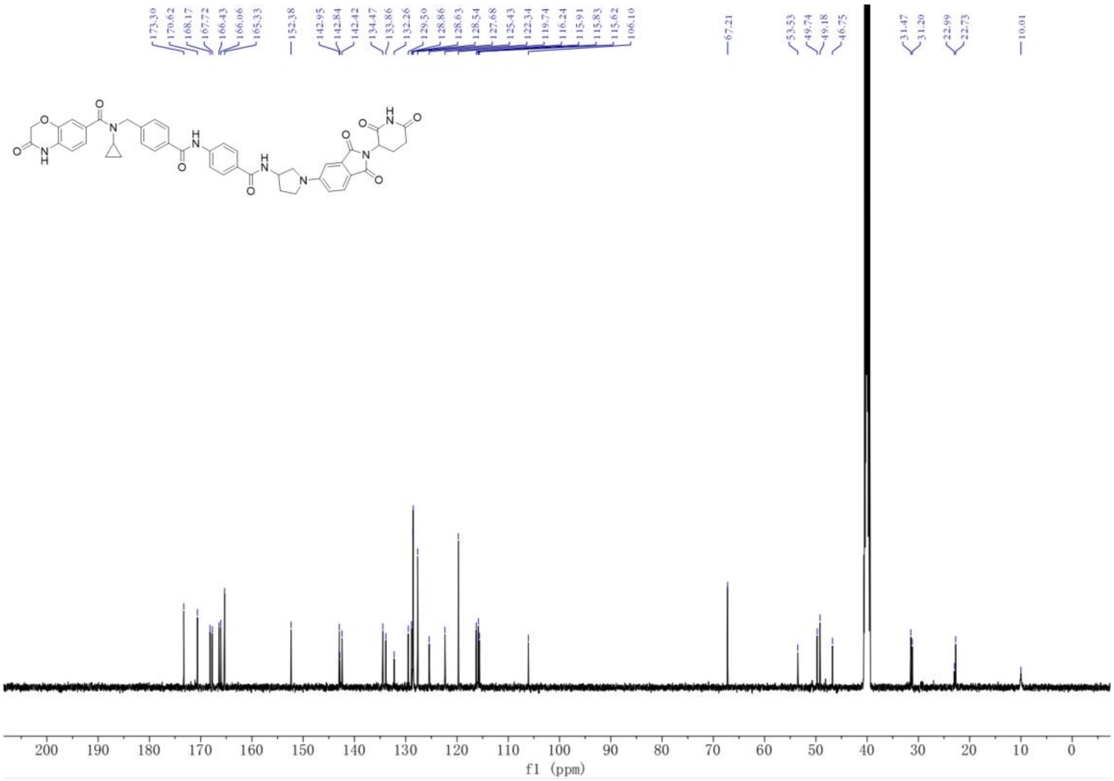

[1]H NMR and [13]C NMR spectra for **8b**

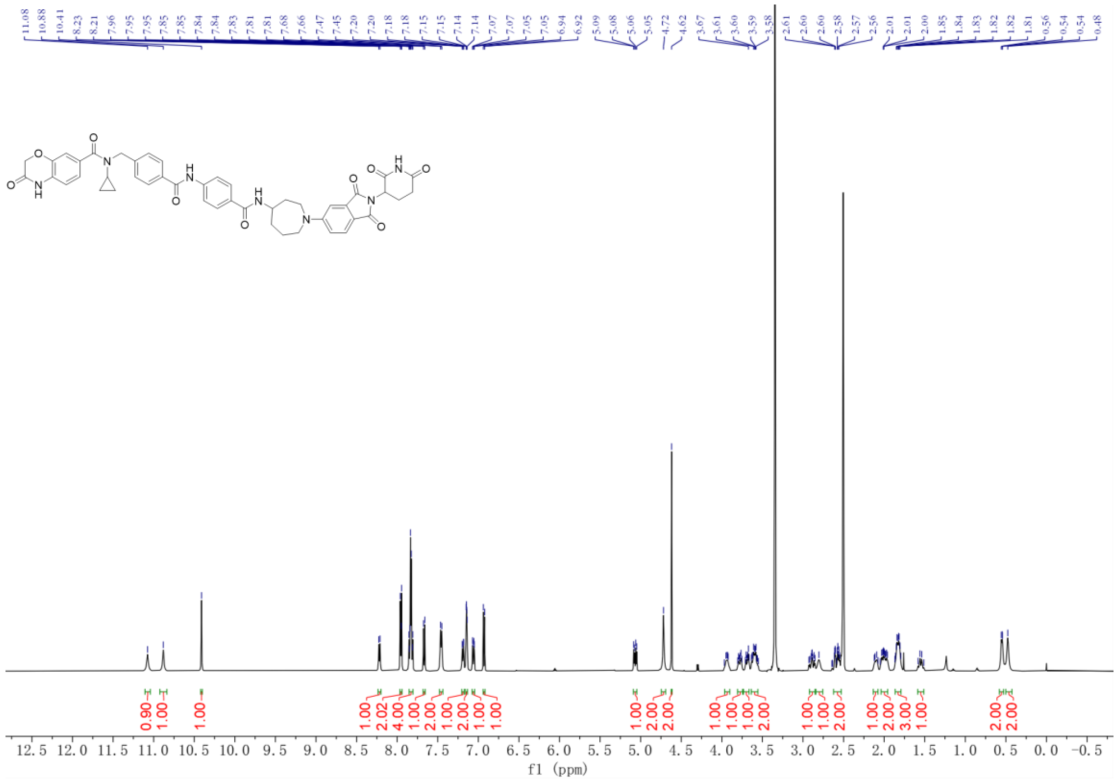


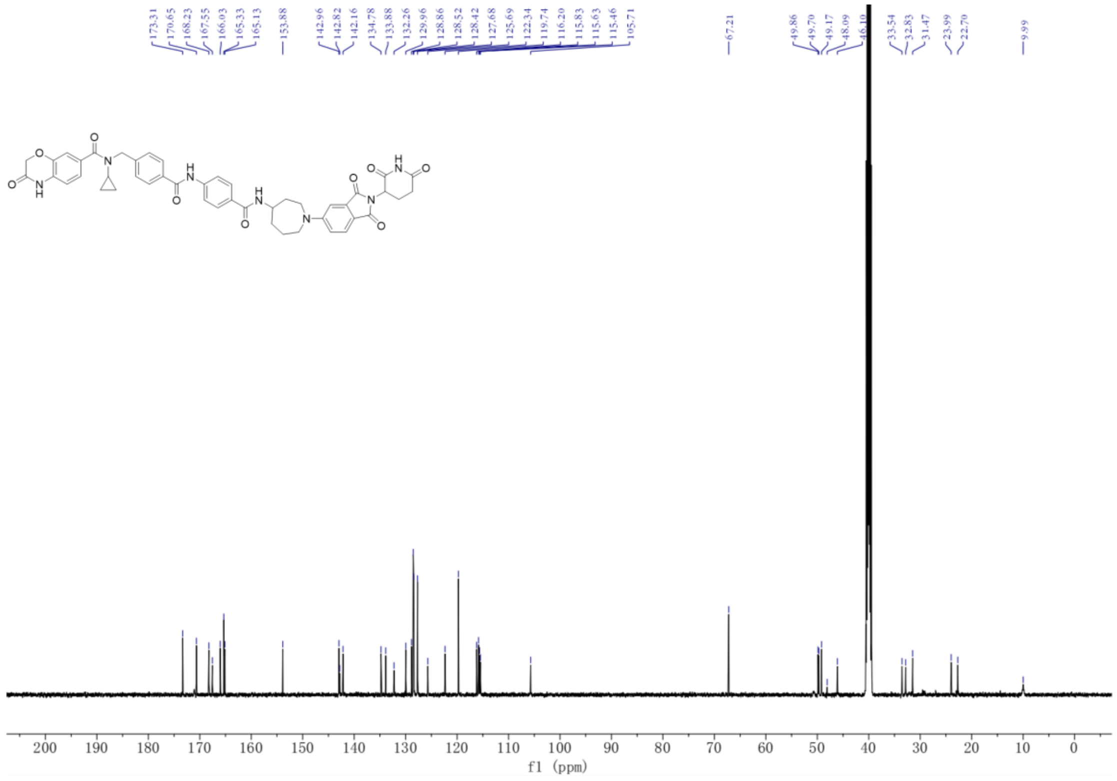

[1]H NMR and [13]C NMR spectra for **8c**

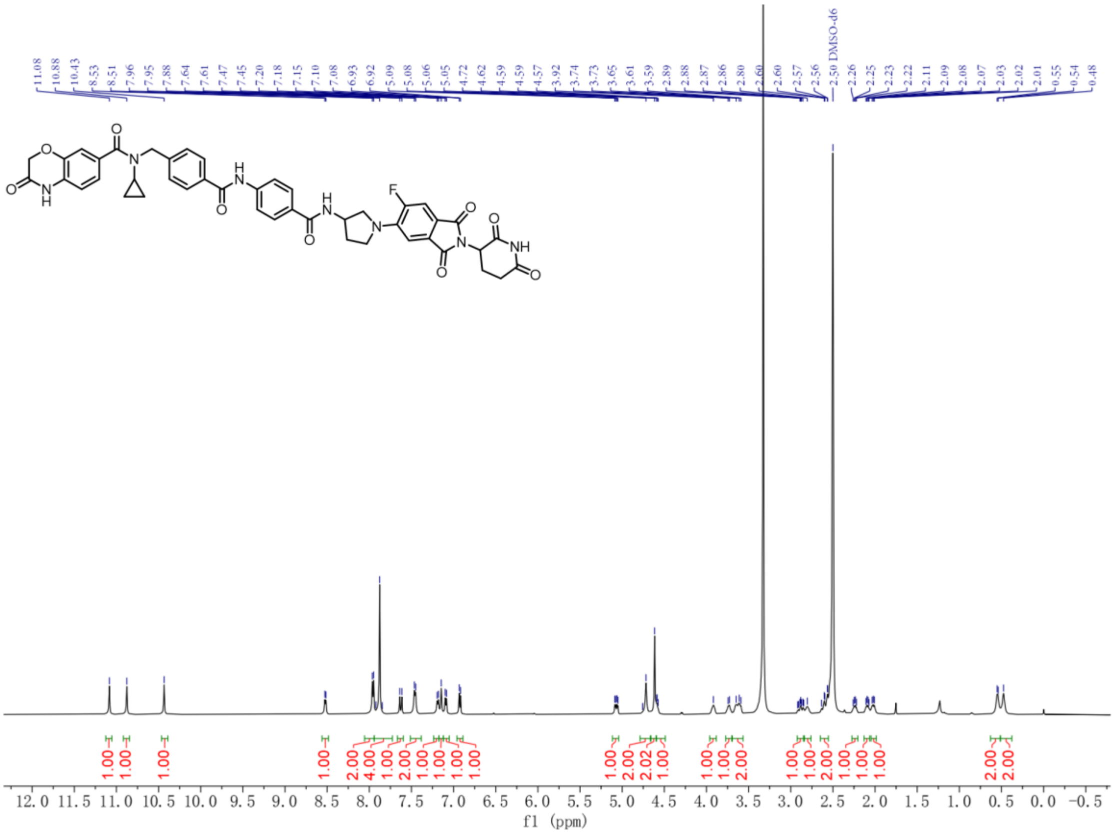


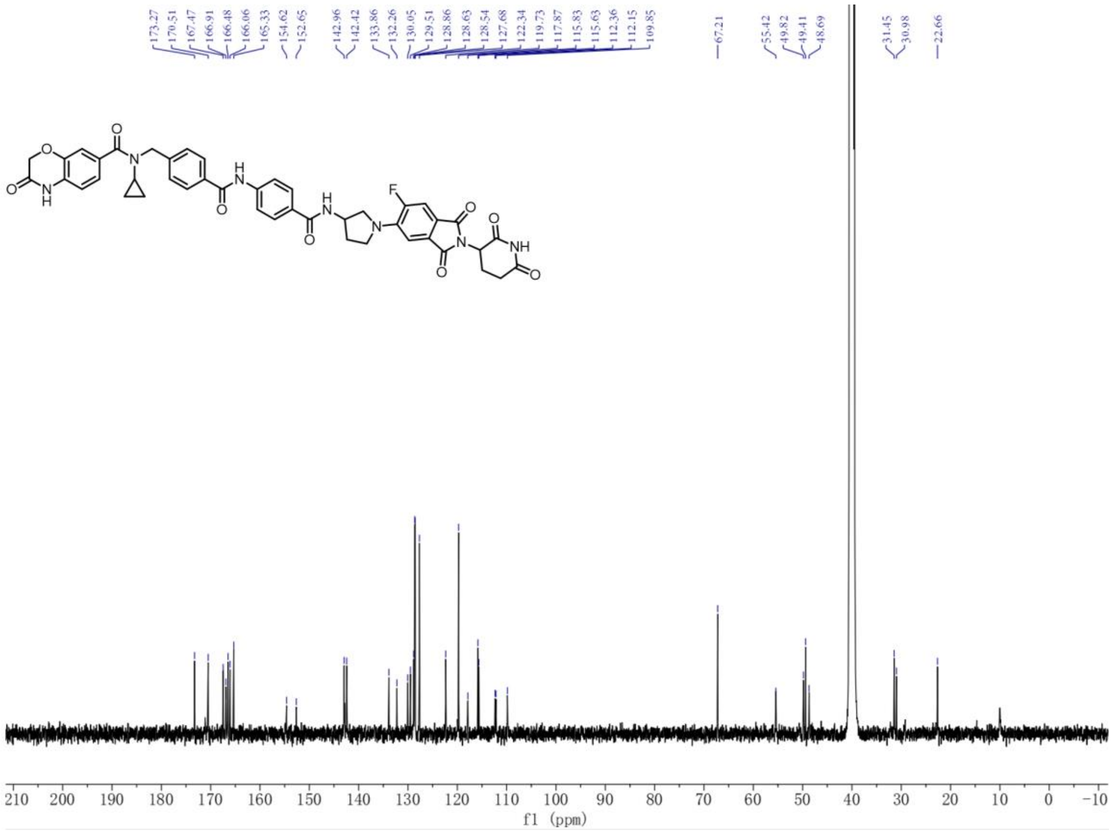

$^{1}H$ NMR and $^{13}C$ NMR spectra for **8d**

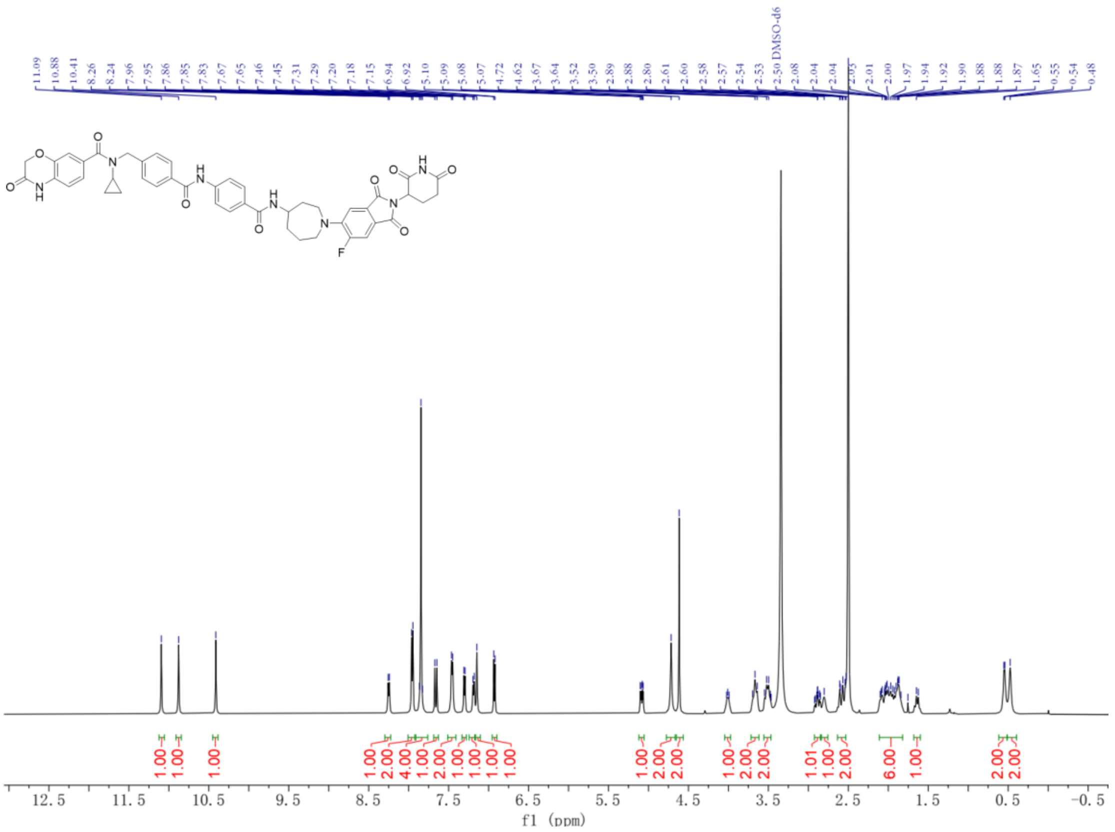


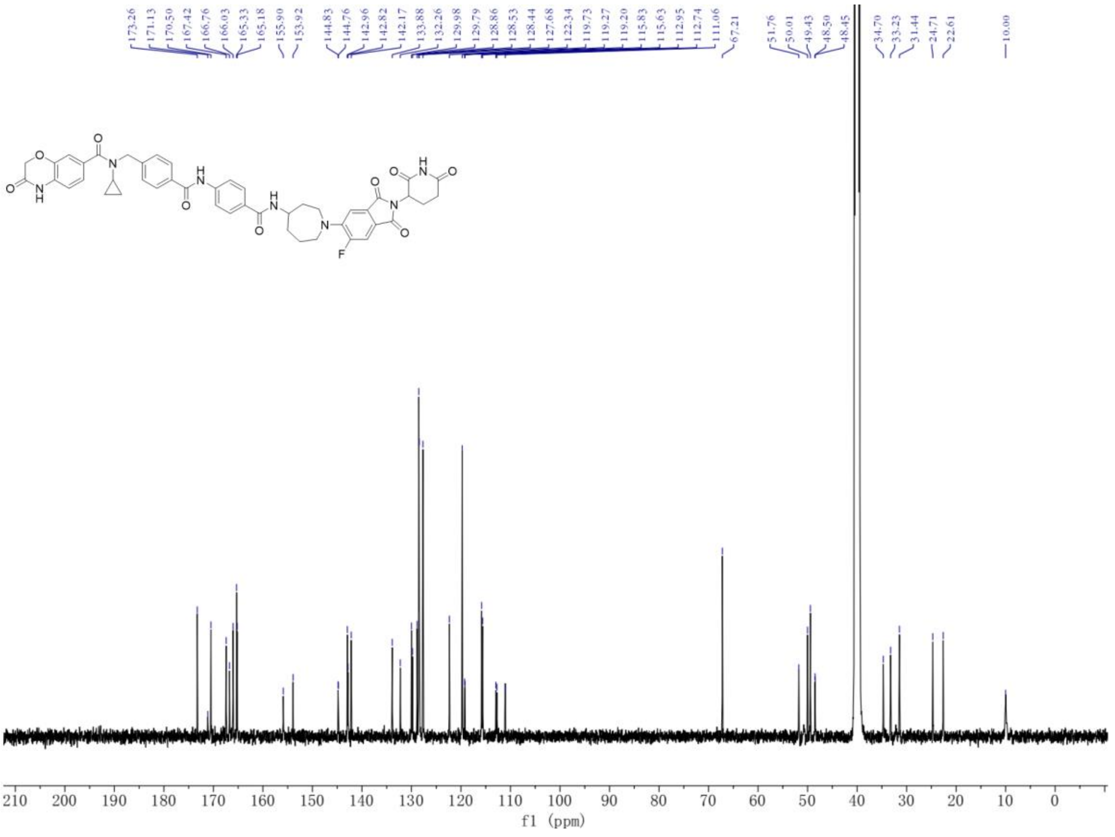

$^{1}$H NMR and $^{13}$C NMR spectra for **8e (424-ND)**

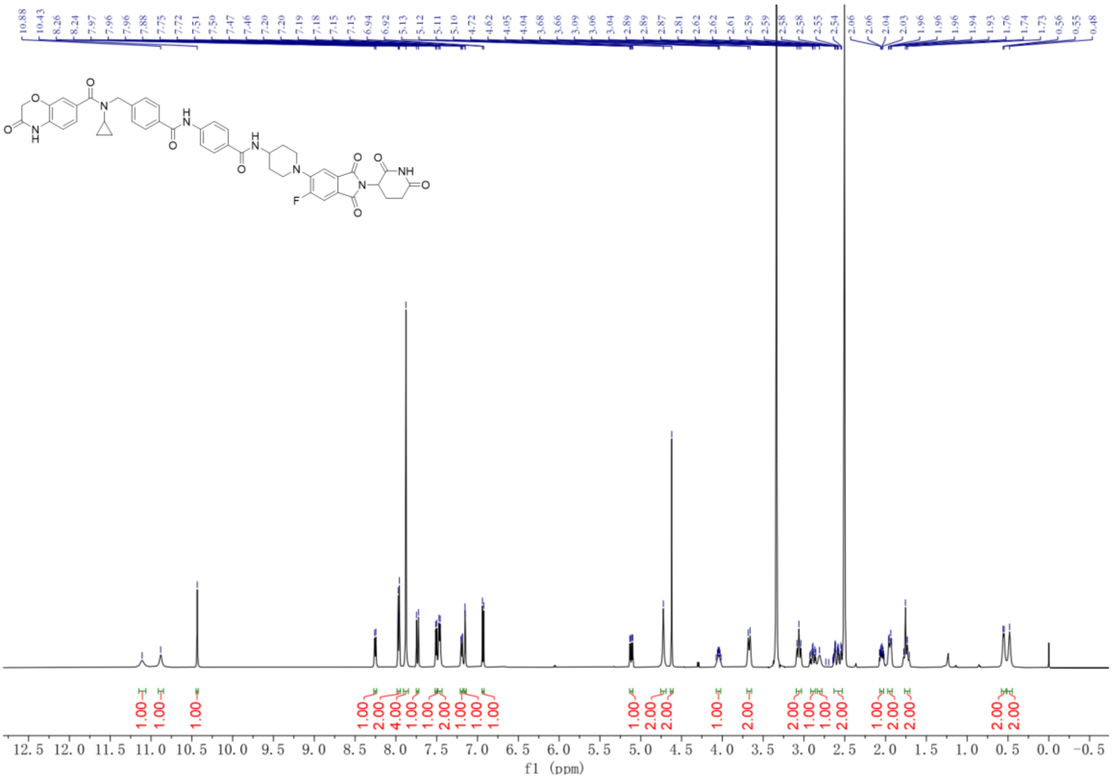


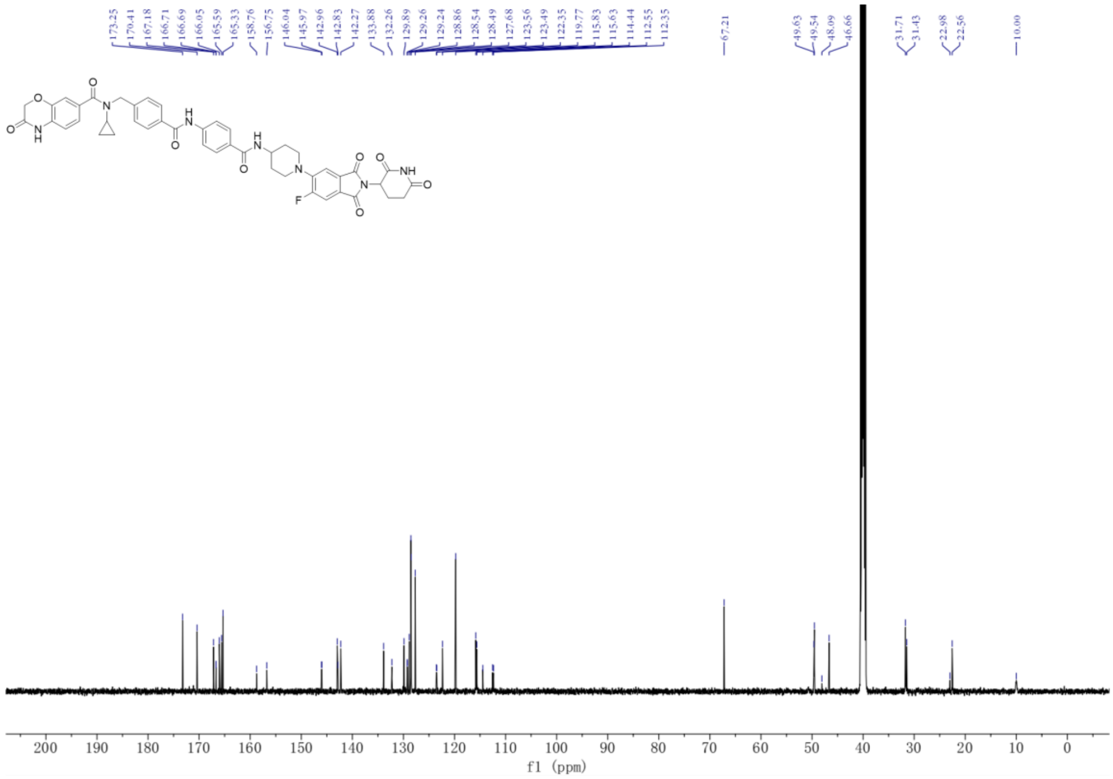

[1]H NMR and [13]C NMR spectra for **8b-S**

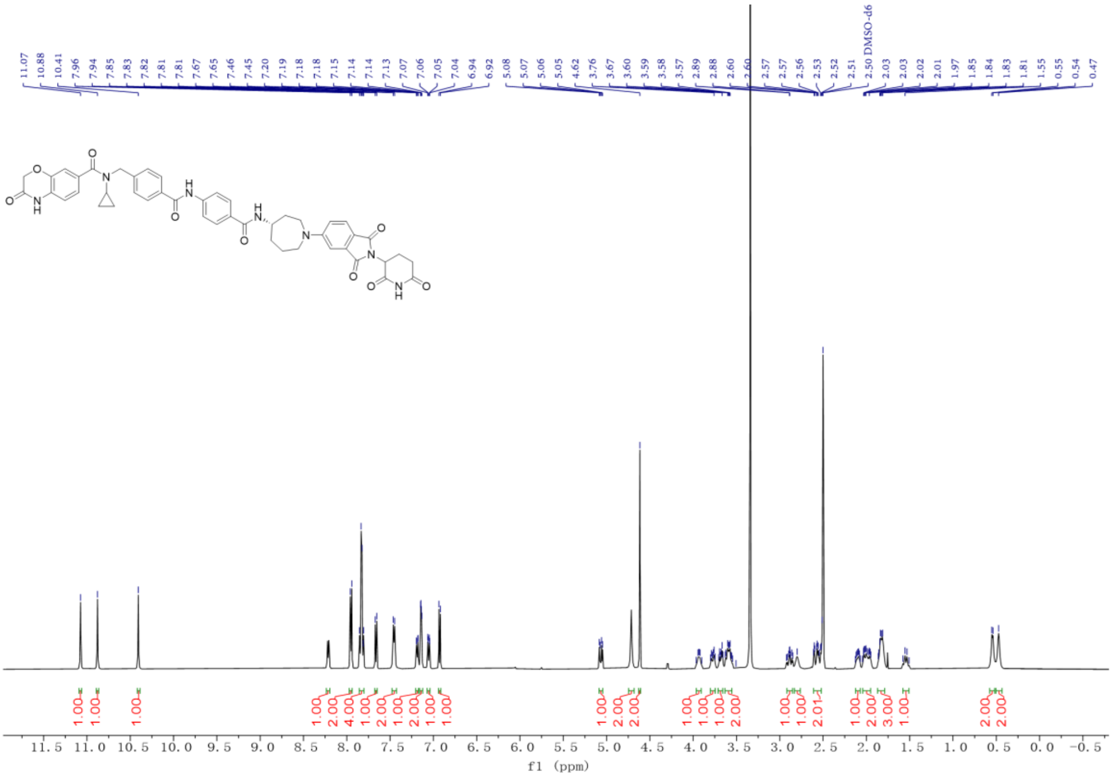


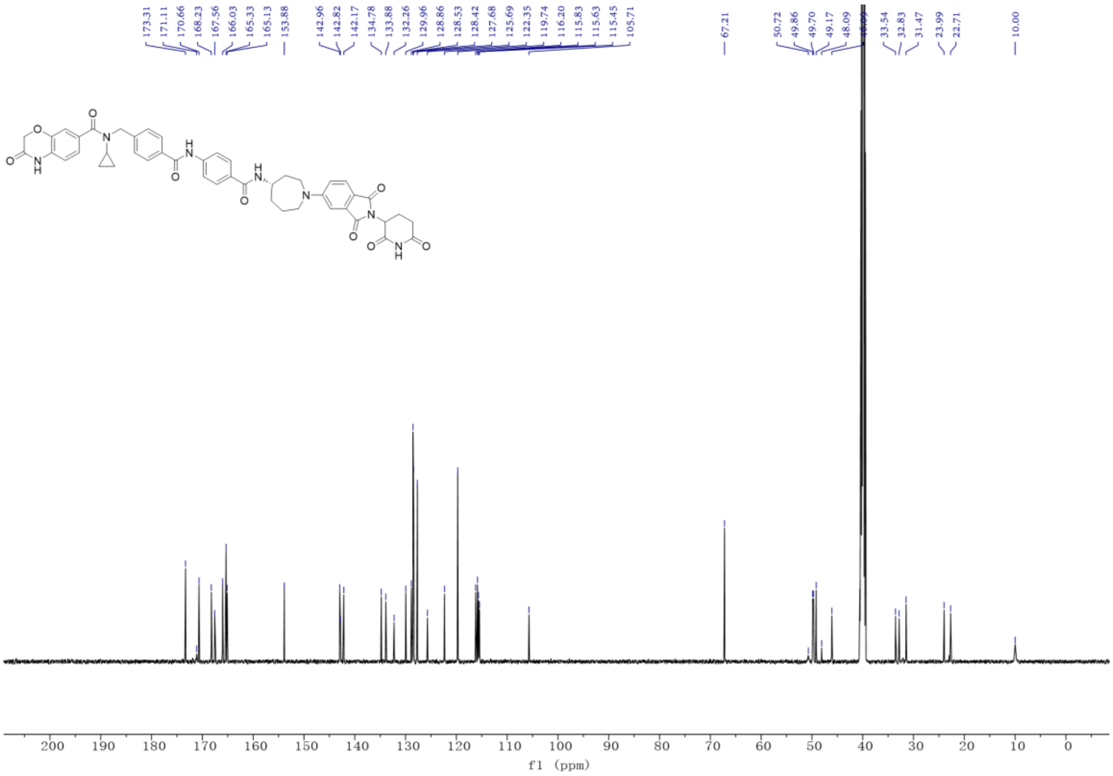

[1]H NMR and [13]C NMR spectra for **8b-R(424-GD)**

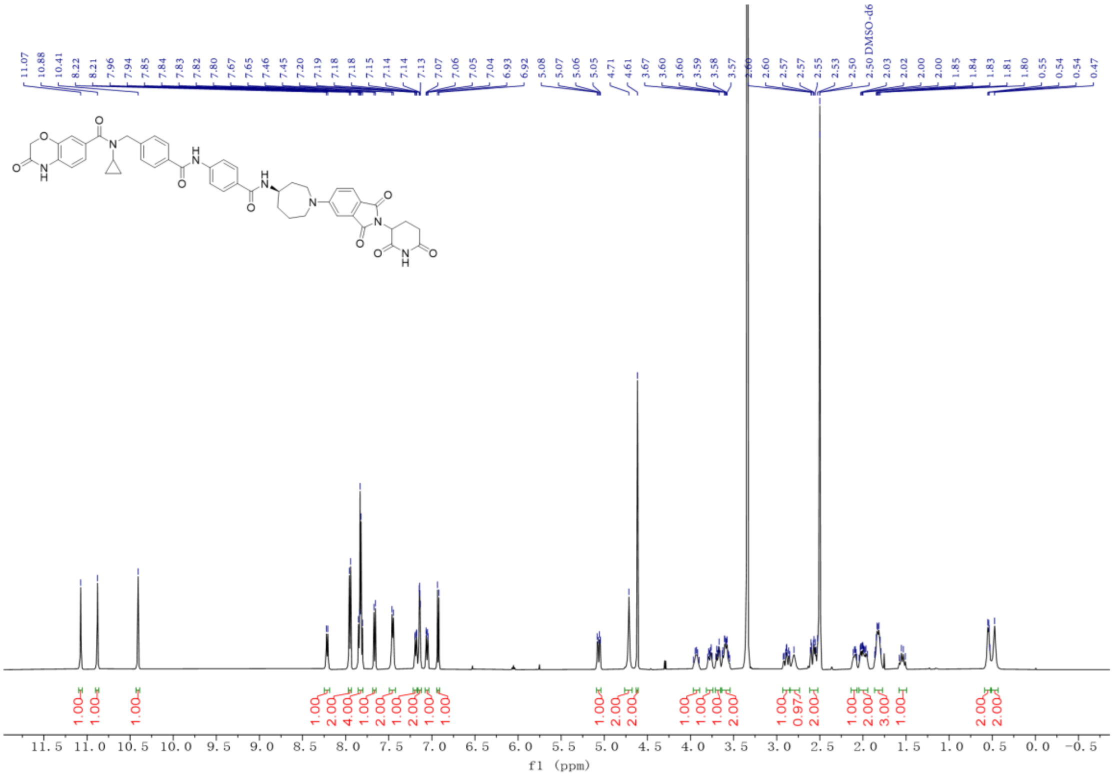


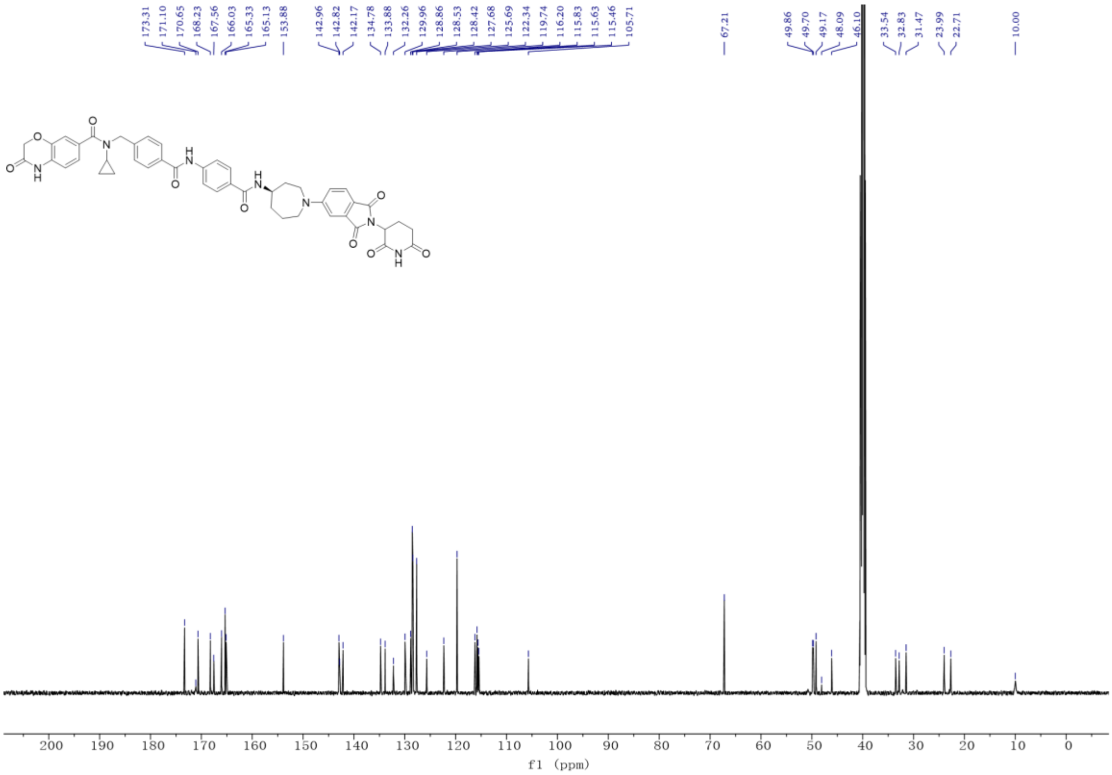

[1]H NMR and [13]C NMR spectra for **424-ND-N1**

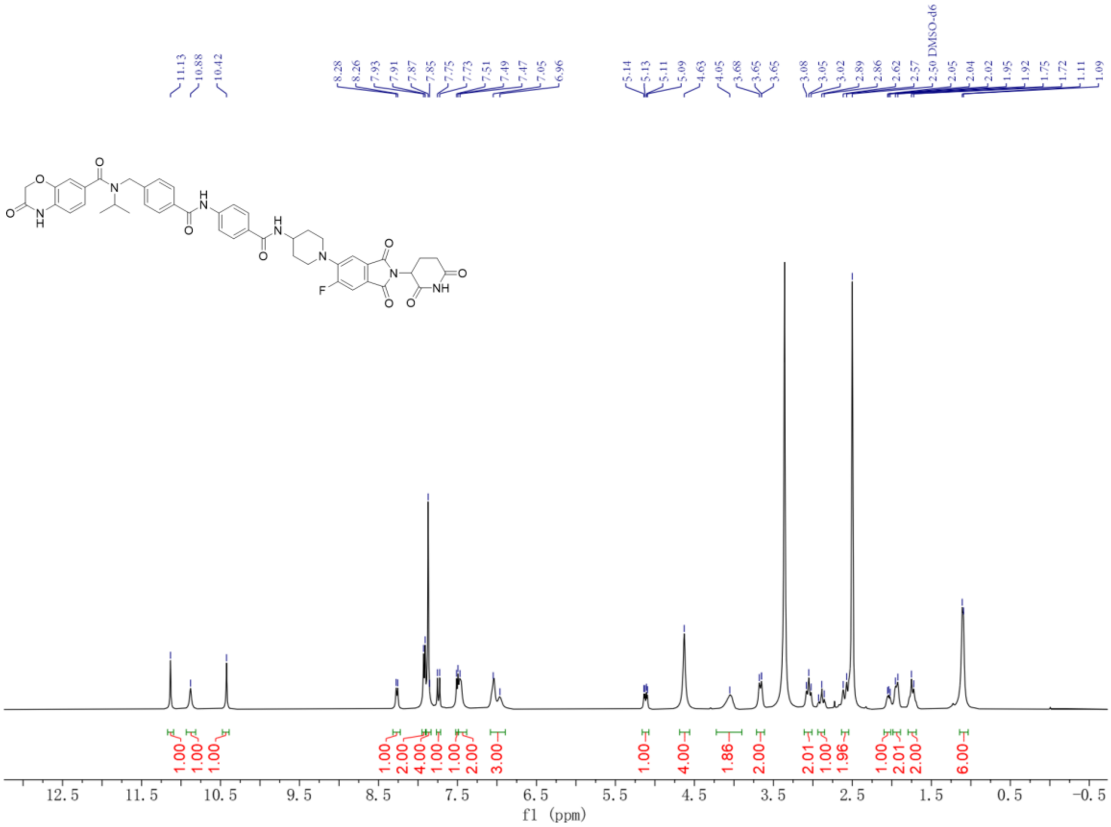


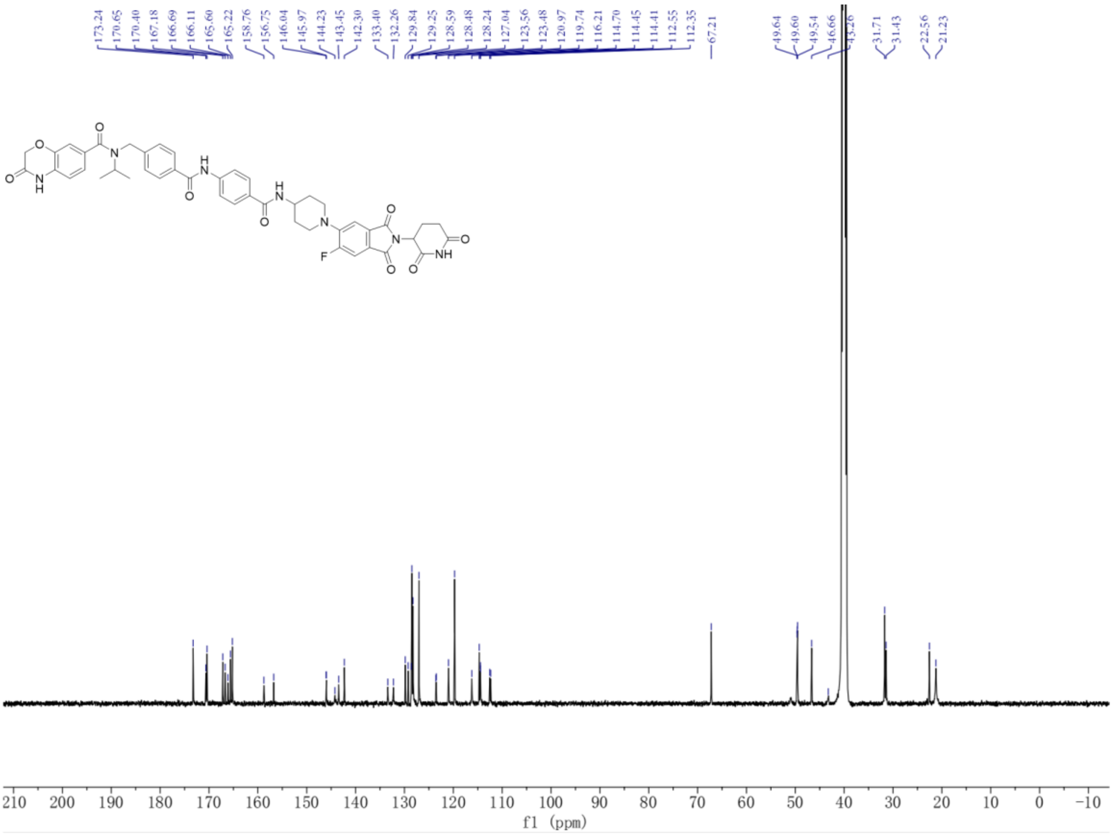

[1]H NMR and [13]C NMR spectra for **424-ND-N2**

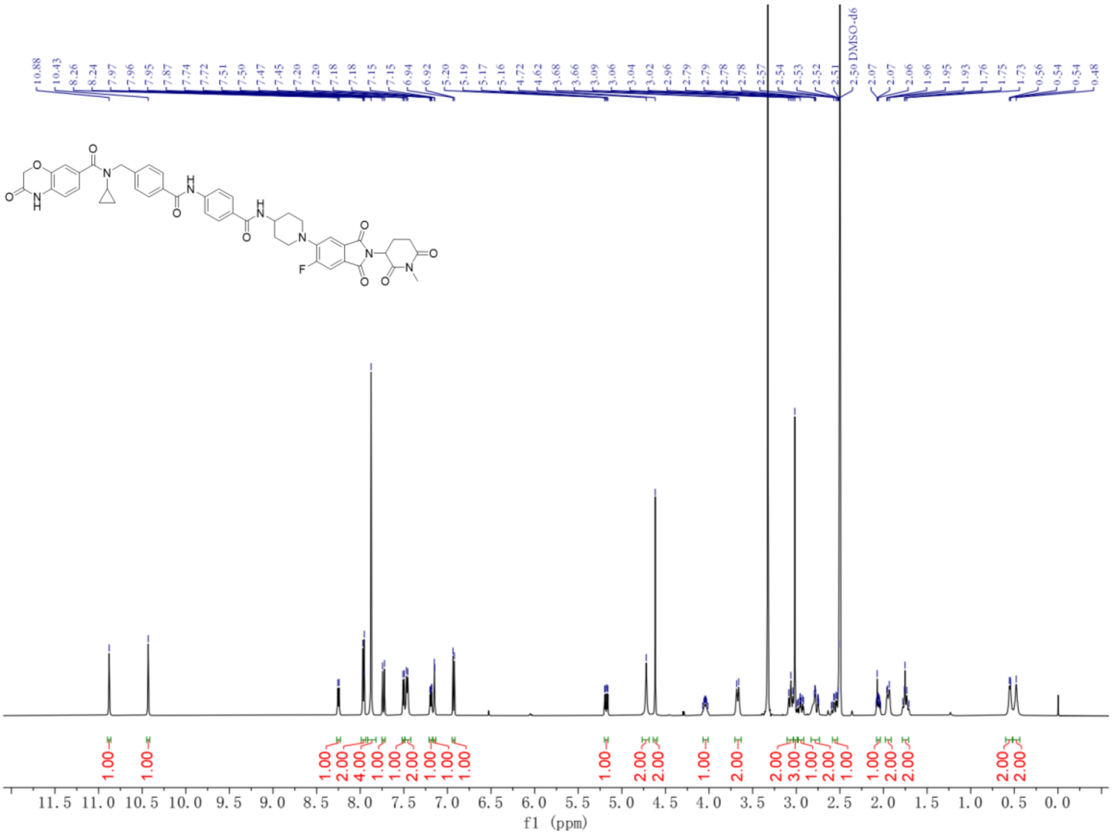

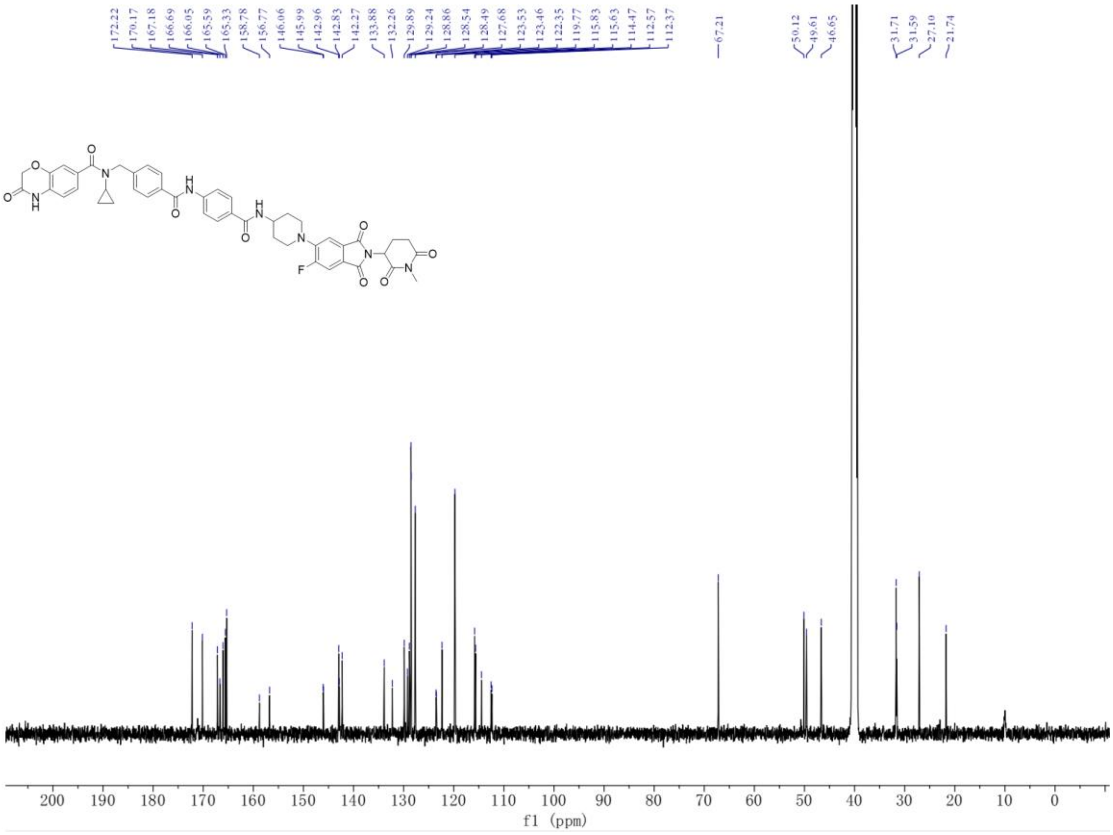

$^{1}$H NMR and $^{13}$C NMR spectra for **424-GD-N1**

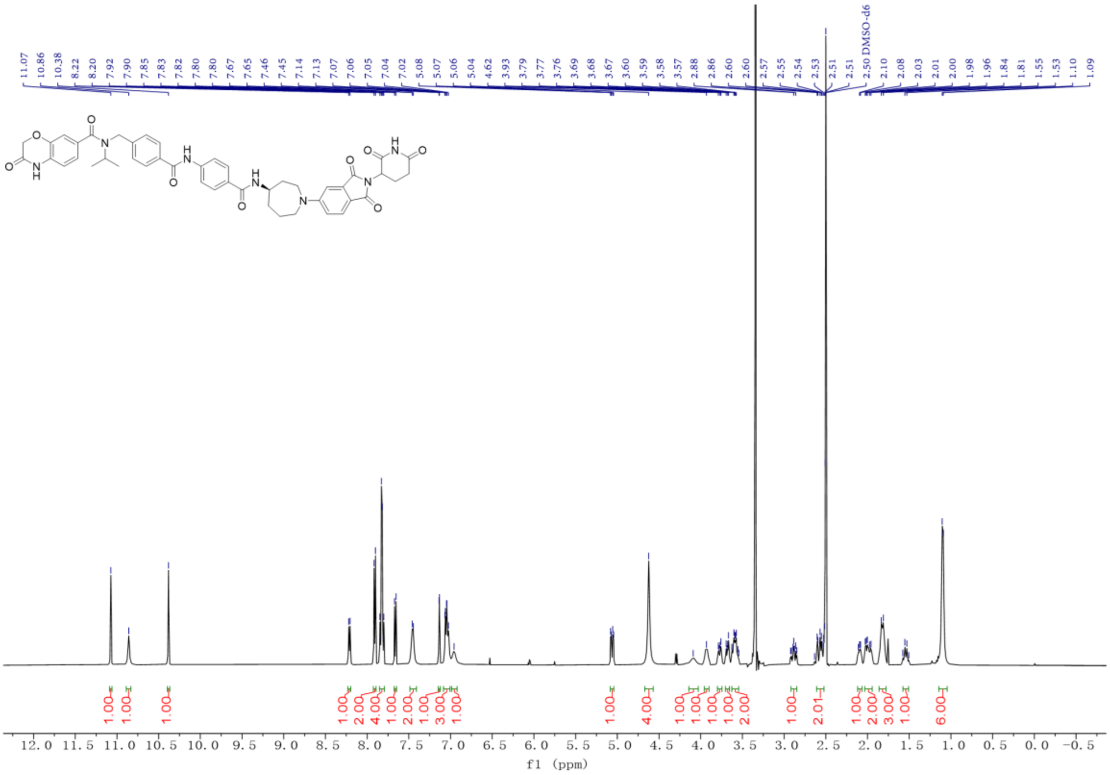


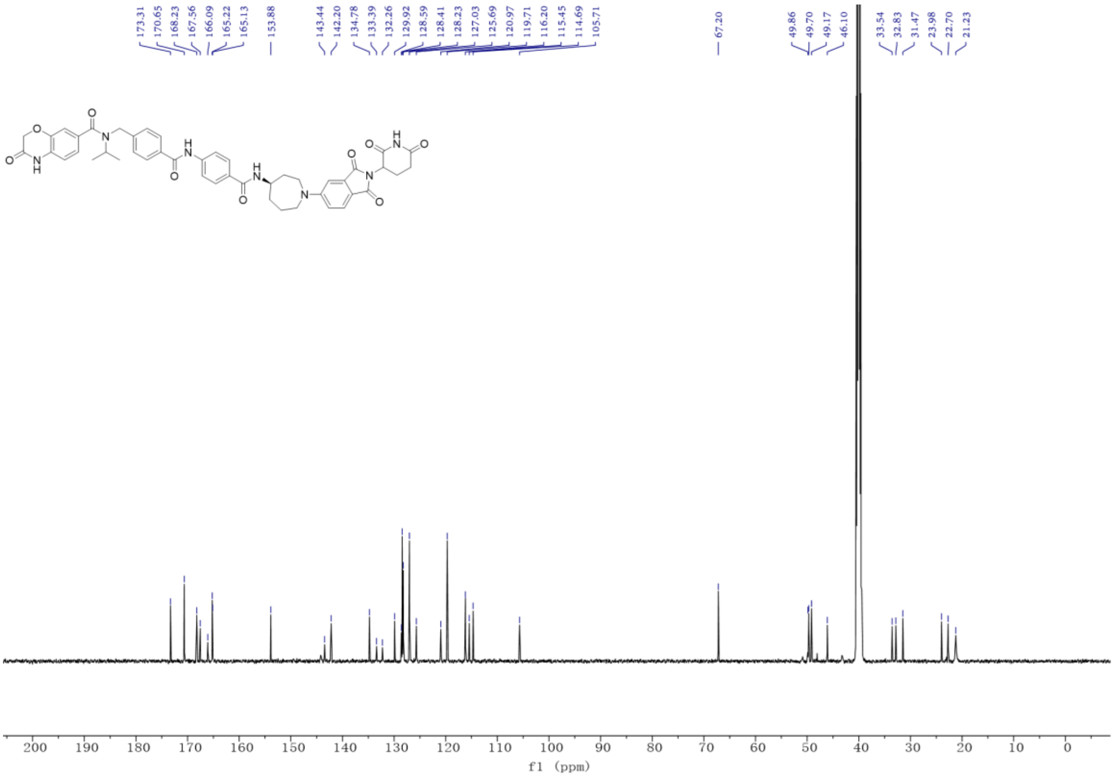

[1]H NMR and [13]C NMR spectra for **424-GD-N2**

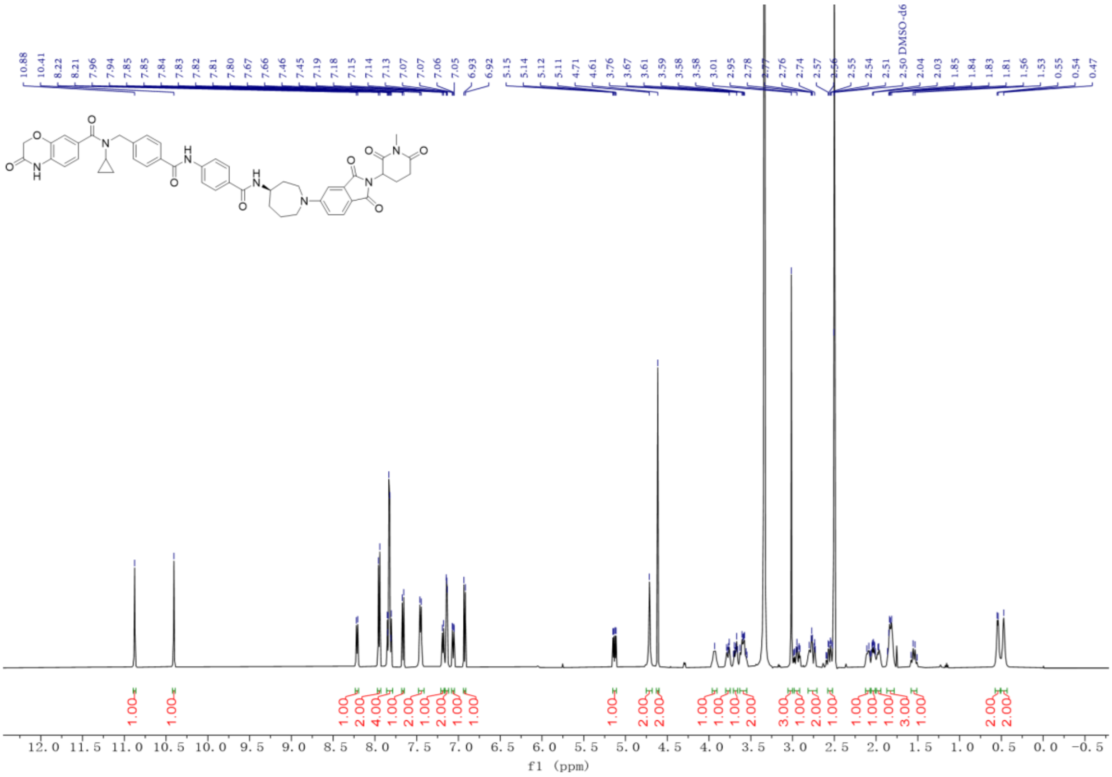


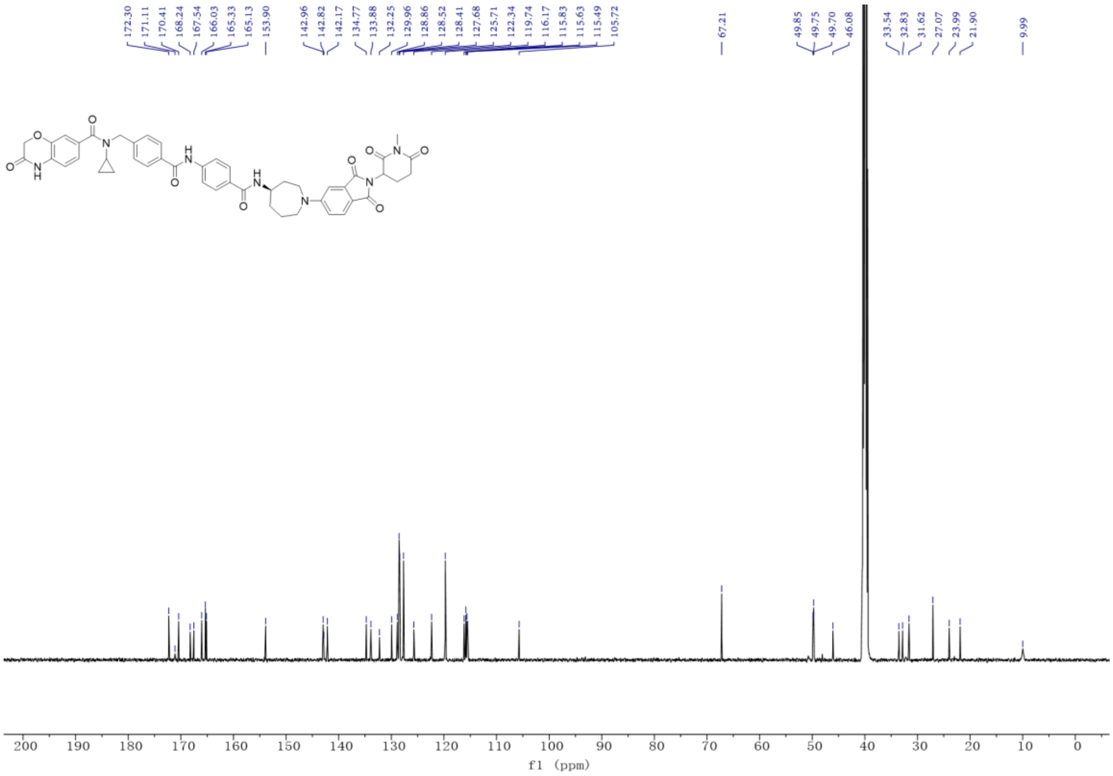

### 8. HPLC spectra of final compounds

HPLC Purity Analysis of **8a**

| | | | |
|---|---|---|---|
| **Data file:** | LLC-5-64-3.dx | | |
| **Sequence Name:** | SingleSample | **Project Name:** | 1260 |
| **Sample name:** | LLC-5-64-3 | **Operator:** | SYSTEM |
| **Instrument:** | 1260 | **Injection date:** | 2023-04-24 21:05:01+08:00 |
| **Inj. volume:** | 10.000 µL | **Location:** | P1-E1 |
| **Acq. method:** | normal.amx | **Type:** | Sample |
| **Processing method:** | INM.pmx | **Sample amount:** | 0.00 |
| **Manually modified:** | Manual Integration | | |

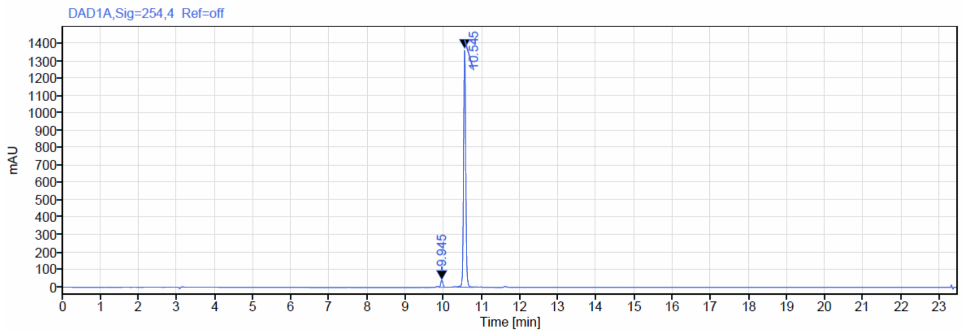


**Signal:** DAD1A,Sig=254,4 Ref=off

| RT [min] | Type | Width [min] | Area | Height | Area% | Name |
|---|---|---|---|---|---|---|
| 9.945 | VB | 0.19 | 157.85 | 40.90 | 2.78 | |
| 10.545 | VV | 0.43 | 5513.31 | 1369.42 | 97.22 | |
| | | **Sum** | 5671.16 | | 100.00 | |

HPLC Purity Analysis of **8b**

| | | | |
|---|---|---|---|
| **Data file:** | LLC-6-74.dx | | |
| **Sequence Name:** | SingleSample | **Project Name:** | 1260 |
| **Sample name:** | LLC-6-74 | **Operator:** | SYSTEM |
| **Instrument:** | 1260 | **Injection date:** | 2023-06-04 14:16:00+08:00 |
| **Inj. volume:** | 20.000 µL | **Location:** | P1-D1 |
| **Acq. method:** | LC-NORMAL-D-B.amx | **Type:** | Sample |
| **Processing method:** | INM.pmx | **Sample amount:** | 0.00 |
| **Manually modified:** | Manual Integration | | |

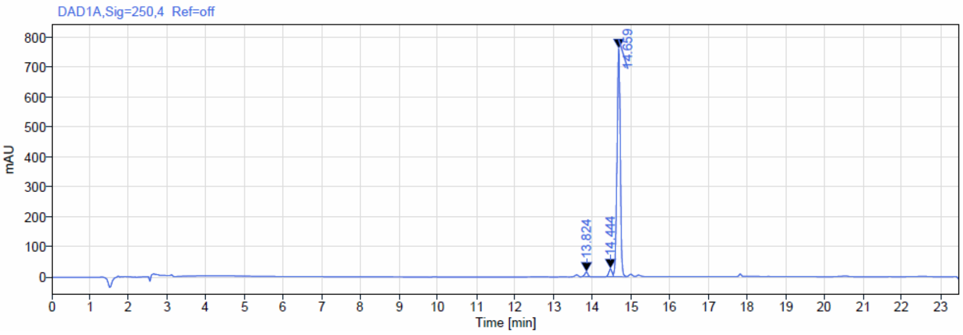


**Signal:** DAD1A,Sig=250,4 Ref=off

| RT [min] | Type | Width [min] | Area | Height | Area% | Name |
|---|---|---|---|---|---|---|
| 13.824 | MM m | 0.20 | 80.08 | 14.74 | 1.70 | |
| 14.444 | MM m | 0.17 | 139.47 | 25.50 | 2.96 | |
| 14.659 | MV m | 0.35 | 4498.05 | 760.90 | 95.35 | |
| | | **Sum** | 4717.60 | | 100.00 | |

HPLC Purity Analysis of **8c**

| | | | |
|---|---|---|---|
| **Data file:** | swz-11-061.dx | | |
| **Sequence Name:** | SingleSample | **Project Name:** | 1260 |
| **Sample name:** | swz-11-061 | **Operator:** | SYSTEM |
| **Instrument:** | 1260 | **Injection date:** | 2025-04-08 18:28:55+07:00 |
| **Inj. volume:** | 10.000 µL | **Location:** | P1-C3 |
| **Acq. method:** | normal.amx | **Type:** | Sample |
| **Processing method:** | INM.pmx | **Sample amount:** | 0.00 |
| **Manually modified:** | Manual Integration | | |

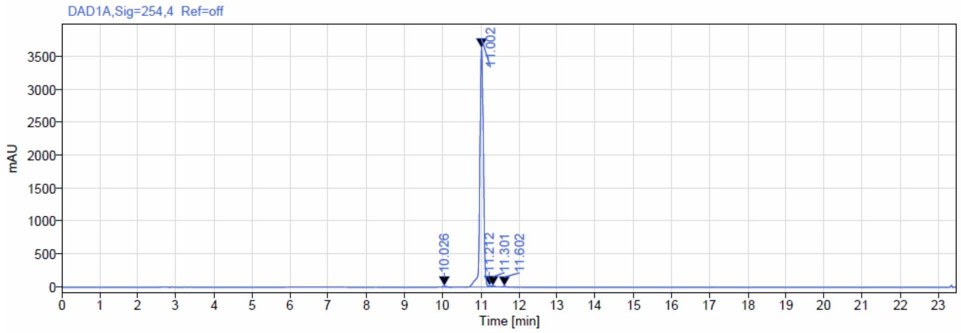


**Signal:** DAD1A,Sig=254,4 Ref=off

| RT [min] | Type | Width [min] | Area | Height | Area% | Name |
|---|---|---|---|---|---|---|
| 10.026 | VB | 0.30 | 131.10 | 16.05 | 0.55 | |
| 11.002 | BV | 0.61 | 23537.49 | 3631.20 | 98.43 | |
| 11.212 | VV | 0.10 | 93.82 | 18.33 | 0.39 | |
| 11.301 | VB | 0.18 | 79.72 | 13.96 | 0.33 | |
| 11.602 | BV | 0.26 | 71.51 | 11.27 | 0.30 | |
| | | **Sum** | 23913.64 | | 100.00 | |

HPLC Purity Analysis of **8d**

| | | | |
|---|---|---|---|
| **Data file:** | swz-11-060.dx | | |
| **Sequence Name:** | SingleSample | **Project Name:** | 1260 |
| **Sample name:** | swz-11-060 | **Operator:** | SYSTEM |
| **Instrument:** | 1260 | **Injection date:** | 2025-04-08 18:03:50+07:00 |
| **Inj. volume:** | 10.000 µL | **Location:** | P1-C2 |
| **Acq. method:** | normal.amx | **Type:** | Sample |
| **Processing method:** | INM.pmx | **Sample amount:** | 0.00 |
| **Manually modified:** | Manual Integration | | |

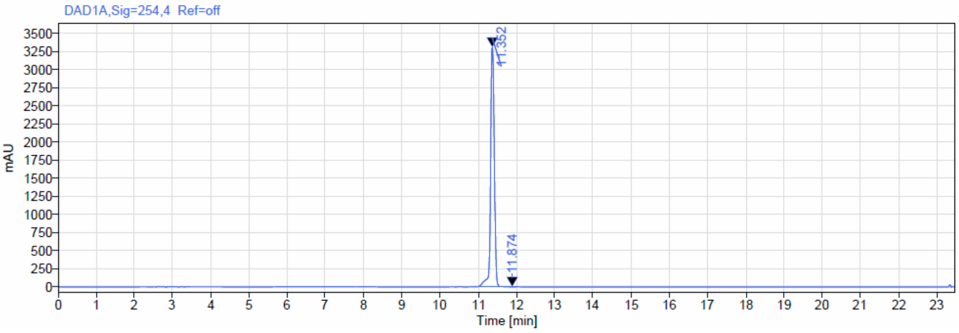


**Signal:** DAD1A,Sig=254,4 Ref=off

| RT [min] | Type | Width [min] | Area | Height | Area% | Name |
|---|---|---|---|---|---|---|
| 11.352 | VV | 0.88 | 22616.87 | 3315.04 | 99.94 | |
| 11.874 | VV | 0.18 | 14.40 | 1.92 | 0.06 | |
| | | **Sum** | 22631.27 | | 100.00 | |

HPLC Purity Analysis of **8e (424-ND)**

| | | | |
|---|---|---|---|
| **Data file:** | LLC-5-48.dx | | |
| **Sequence Name:** | SingleSample | **Project Name:** | 1260 |
| **Sample name:** | LLC-5-48 | **Operator:** | SYSTEM |
| **Instrument:** | 1260 | **Injection date:** | 2023-03-10 11:34:57+08:00 |
| **Inj. volume:** | 15.000 µL | **Location:** | P1-A1 |
| **Acq. method:** | normal.amx | **Type:** | Sample |
| **Processing method:** | GC_LC 面积百分比_DefaultMethod.pmx | **Sample amount:** | 0.00 |
| **Manually modified:** | Manual Integration | | |

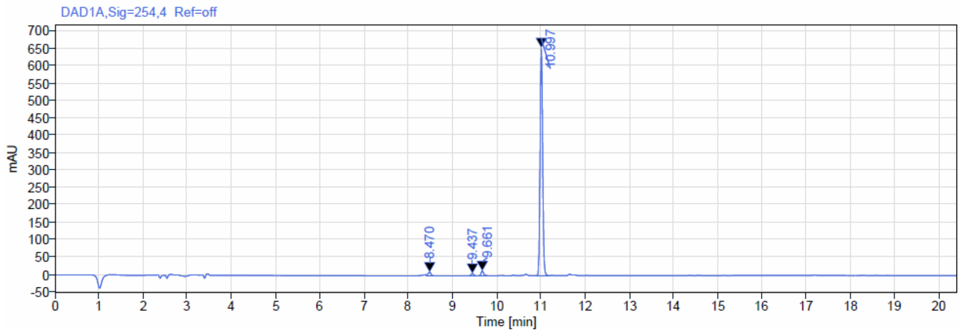


**Signal:** DAD1A,Sig=254,4 Ref=off

| RT [min] | Type | Width [min] | Area | Height | Area% | Name |
|---|---|---|---|---|---|---|
| 8.470 | VB | 0.27 | 50.42 | 10.48 | 1.82 | |
| 9.437 | VB | 0.21 | 25.24 | 6.57 | 0.91 | |
| 9.661 | BV | 0.29 | 60.22 | 13.90 | 2.17 | |
| 10.997 | BV | 0.31 | 2635.85 | 654.68 | 95.10 | |
| | | **Sum** | 2771.74 | | 100.00 | |

HPLC Purity Analysis of **8b-S**

| | | | |
|---|---|---|---|
| **Data file:** | swz-16-80.dx | | |
| **Sequence Name:** | SingleSample | **Project Name:** | 1260 |
| **Sample name:** | swz-16-80 | **Operator:** | SYSTEM |
| **Instrument:** | 1260 | **Injection date:** | 2026-03-06 10:23:11+07:00 |
| **Inj. volume:** | 5.000 µL | **Location:** | P1-E5 |
| **Acq. method:** | normal.amx | **Type:** | Sample |
| **Processing method:** | INM.pmx | **Sample amount:** | 0.00 |
| **Manually modified:** | Manual Integration | | |

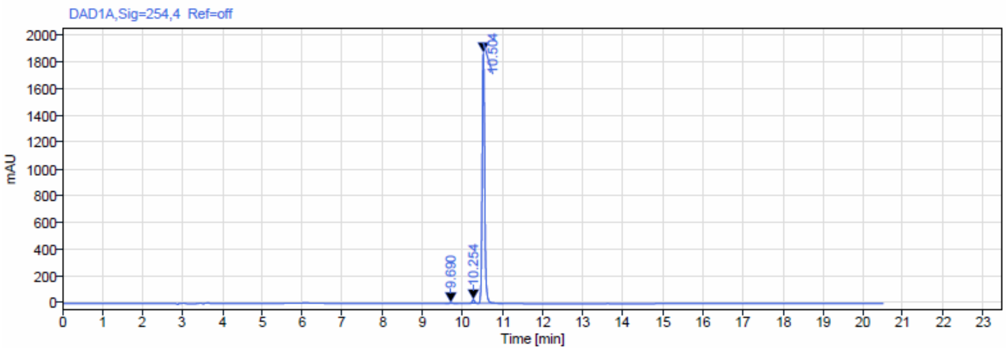


**Signal:** DAD1A,Sig=254,4 Ref=off

| RT [min] | Type | Width [min] | Area | Height | Area% | Name |
|---|---|---|---|---|---|---|
| 9.690 | BV | 0.32 | 38.68 | 6.54 | 0.46 | |
| 10.254 | VV | 0.20 | 126.46 | 29.46 | 1.50 | |
| 10.504 | VV | 0.63 | 8255.37 | 1874.04 | 98.04 | |
| | | **Sum** | 8420.51 | | 100.00 | |

HPLC Purity Analysis of **8b-R(424-GD)**

| | | | |
|---|---|---|---|
| **Data file:** | swz-16-81.dx | | |
| **Sequence Name:** | SingleSample | **Project Name:** | 1260 |
| **Sample name:** | swz-16-81 | **Operator:** | SYSTEM |
| **Instrument:** | 1260 | **Injection date:** | 2026-03-06 09:56:38+07:00 |
| **Inj. volume:** | 5.000 µL | **Location:** | P1-E4 |
| **Acq. method:** | normal.amx | **Type:** | Sample |
| **Processing method:** | INM.pmx | **Sample amount:** | 0.00 |
| **Manually modified:** | Manual Integration | | |

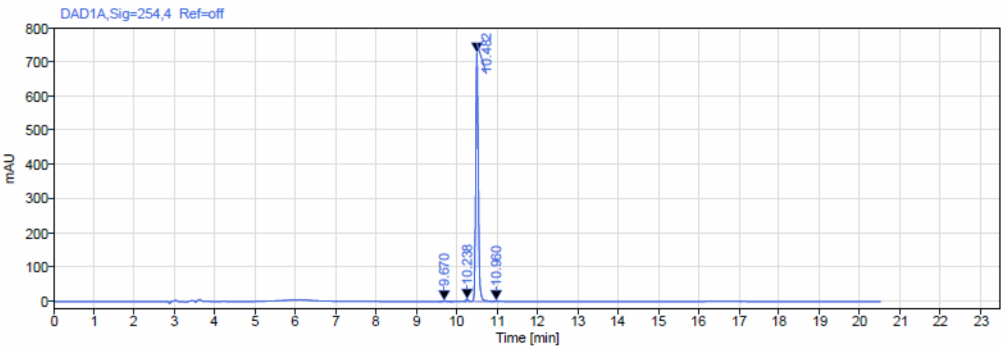


**Signal:** DAD1A,Sig=254,4 Ref=off

| RT [min] | Type | Width [min] | Area | Height | Area% | Name |
|---|---|---|---|---|---|---|
| 9.670 | BV | 0.35 | 15.30 | 2.60 | 0.47 | |
| 10.238 | VB | 0.18 | 28.35 | 7.05 | 0.87 | |
| 10.482 | BV | 0.55 | 3192.62 | 726.67 | 98.24 | |
| 10.960 | VV | 0.14 | 13.58 | 2.88 | 0.42 | |
| | | **Sum** | 3249.85 | | 100.00 | |

HPLC Purity Analysis of **424-ND-N1**

| | | | |
|---|---|---|---|
| **Data file:** | swz-11-063.dx | | |
| **Sequence Name:** | SingleSample | **Project Name:** | 1260 |
| **Sample name:** | swz-11-063 | **Operator:** | SYSTEM |
| **Instrument:** | 1260 | **Injection date:** | 2025-04-09 19:54:37+07:00 |
| **Inj. volume:** | 10.000 µL | **Location:** | P1-B2 |
| **Acq. method:** | normal.amx | **Type:** | Sample |
| **Processing method:** | INM.pmx | **Sample amount:** | 0.00 |
| **Manually modified:** | Manual Integration | | |

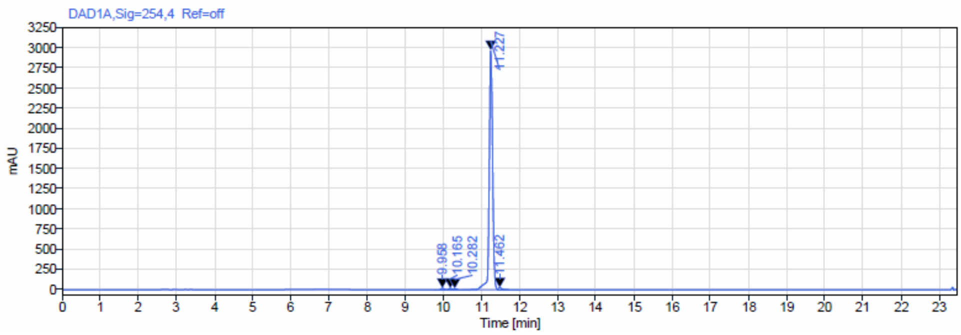


**Signal:** DAD1A,Sig=254,4 Ref=off

| RT [min] | Type | Width [min] | Area | Height | Area% | Name |
|---|---|---|---|---|---|---|
| 9.958 | BV | 0.37 | 103.51 | 18.73 | 0.55 | |
| 10.165 | VV | 0.15 | 89.51 | 17.10 | 0.48 | |
| 10.282 | VB | 0.25 | 70.23 | 12.80 | 0.38 | |
| 11.227 | BV | 0.92 | 18244.48 | 2975.93 | 97.70 | |
| 11.462 | VB | 0.58 | 166.15 | 24.10 | 0.89 | |
| | | **Sum** | 18673.89 | | 100.00 | |

HPLC Purity Analysis of **424-GD-N1**

| | | | |
|---|---|---|---|
| **Data file:** | swz-16-85.dx | | |
| **Sequence Name:** | SingleSample | **Project Name:** | 1260 |
| **Sample name:** | swz-16-85 | **Operator:** | SYSTEM |
| **Instrument:** | 1260 | **Injection date:** | 2026-03-06 11:16:15+07:00 |
| **Inj. volume:** | 5.000 µL | **Location:** | P1-E7 |
| **Acq. method:** | normal.amx | **Type:** | Sample |
| **Processing method:** | INM.pmx | **Sample amount:** | 0.00 |
| **Manually modified:** | Manual Integration | | |

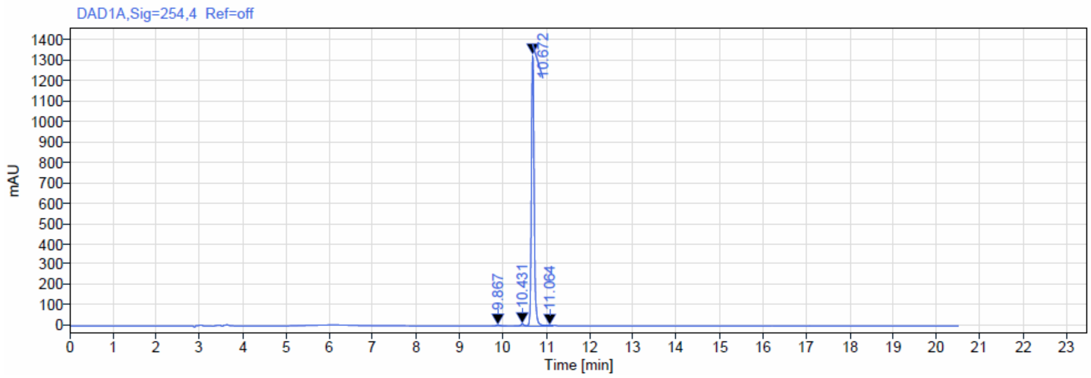


**Signal:** DAD1A,Sig=254,4 Ref=off

| RT [min] | Type | Width [min] | Area | Height | Area% | Name |
|---|---|---|---|---|---|---|
| 9.867 | BB | 0.34 | 23.99 | 4.22 | 0.40 | |
| 10.431 | BV | 0.28 | 45.60 | 10.66 | 0.76 | |
| 10.672 | VV | 0.48 | 5872.60 | 1329.80 | 98.50 | |
| 11.064 | VV | 0.14 | 20.07 | 3.11 | 0.34 | |
| | | **Sum** | 5962.25 | | 100.00 | |

HPLC Purity Analysis of **424-GD-N2**

| | | | |
|---|---|---|---|
| **Data file:** | swz-16-87.dx | | |
| **Sequence Name:** | SingleSample | **Project Name:** | 1260 |
| **Sample name:** | swz-16-87 | **Operator:** | SYSTEM |
| **Instrument:** | 1260 | **Injection date:** | 2026-03-06 00:12:28+07:00 |
| **Inj. volume:** | 8.000 µL | **Location:** | P1-C2 |
| **Acq. method:** | normal.amx | **Type:** | Sample |
| **Processing method:** | INM.pmx | **Sample amount:** | 0.00 |
| **Manually modified:** | Manual Integration | | |

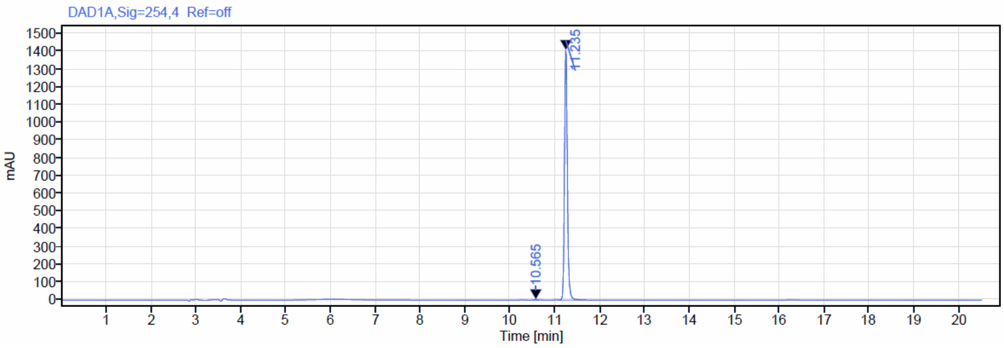


**Signal:** DAD1A,Sig=254,4 Ref=off

| RT [min] | Type | Width [min] | Area | Height | Area% | Name |
|---|---|---|---|---|---|---|
| 10.565 | BV | 0.14 | 11.43 | 2.63 | 0.18 | |
| 11.235 | VM m | 0.60 | 6249.14 | 1407.13 | 99.82 | |
| | | **Sum** | **6260.57** | | | |
